# Dopamine signaling in wake promoting clock neurons is not required for the normal regulation of sleep in *Drosophila*

**DOI:** 10.1101/2020.05.20.106369

**Authors:** Florencia Fernandez-Chiappe, Christiane Hermann-Luibl, Alina Peteranderl, Nils Reinhard, Marie Hieke, Mareike Selcho, Orie T. Shafer, Nara I. Muraro, Charlotte Helfrich-Förster

**Author notes:** these authors contributed equally to this work (Florencia Fernandez-Chiappe, Christiane Hermann-Luibl). Department of Animal Physiology, Institute of Biology, Leipzig University, Leipzig, Germany (Mareike Selcho). To whom all correspondence should be addressed: Charlotte Helfrich-Förster, Neurobiology and Genetics, Theodor Boveri Institute, Biocenter, University of Würzburg, D-97074 Würzburg, Germany.

## Abstract

Dopamine is a wakefulness promoting neuromodulator in mammals and fruit flies. In *D. melanogaster*, the network of clock neurons that drives sleep/activity cycles comprises both wake and sleep promoting cell types, indicating that the sleep-wake circuitry is intimately linked to the circadian clock. The large and small ventrolateral neurons (l-LN_v_s and s-LN_v_s) have been identified as wake-promoting neurons within the clock neuron network. The l-LN_v_s are innervated by dopaminergic neurons, and earlier work proposed that dopamine signaling raises cAMP levels in the l-LN_v_s and thus induces excitatory electrical activity (action potential firing), which results in wakefulness and inhibits sleep. Here, we test this hypothesis by combining cAMP imaging and patch-clamp recordings in isolated brains. We find that dopamine application indeed increases cAMP levels and depolarizes the l-LN_v_s, but surprisingly, it does not result in increased firing rates. Down-regulation of the excitatory dopamine receptor, Dop1R1 in the l-and s-LN_v_s, but not of Dop1R2, abolished the depolarization of l-LN_v_s in response to dopamine. This indicates that dopamine signals via Dop1R1 to the l-LN_v_s. Down-regulation of Dop1R1 or Dop1R2 receptors in the l- and s-LN_v_s does not affect sleep. Unexpectedly, we find a moderate decrease of daytime sleep with down-regulation of Dop1R1 and of nighttime sleep with down-regulation of Dop1R2. Since the l-LN_v_s do not utilize Dop1R2 receptors and the s-LN_v_s respond also to dopamine, we conclude that the s-LN_v_s are responsible for the observed decrease in nighttime sleep. In summary, dopamine signaling in the wake-promoting LN_v_s is not required for daytime arousal, but likely promotes nighttime sleep via the s-LN_v_s.

**Significance statement:** In insect and mammalian brains, sleep promoting networks are intimately linked to the circadian clock, and the mechanisms underlying sleep and circadian timekeeping are evolutionarily ancient and highly conserved. Here we show that dopamine, one important sleep modulator in flies and mammals, plays surprisingly complex roles in the regulation of sleep by clock containing neurons. Dopamine inhibits neurons in a central brain sleep center to promote sleep and excites wake-promoting circadian clock neurons. It is therefore predicted to promote wakefulness through both of these networks. Nevertheless, our results reveal that dopamine acting on wake promoting clock neurons promotes sleep, revealing a previously unappreciated complexity in the dopaminergic control of sleep.

## Introduction

The fruit fly *Drosophila melanogaster* has become a powerful and widely-used model system for sleep research (reviewed by Cirelli, 2009; Dubowy and Sehgal, 2017; Helfrich-Förster, 2018). As in mammals, the sleep-like state of *Drosophila* is associated with reduced sensory responsiveness and reduced brain activity (Nitz et al., 2002; van Swinderen et al., 2004), and is subject to both circadian and homeostatic regulation (Hendricks et al., 2000; Shaw et al., 2000). Furthermore, as in mammals, dopamine and octopamine (the insect functional homolog to noradrenaline) promote arousal in fruit flies (Andretic et al., 2005; Kume et al., 2005; Lima and Miesenböck, 2005; Wu et al., 2008, Lebestky et al., 2009; Crocker et al., 2010; Riemensperger et al., 2011), and GABA promotes sleep (Agosto et al., 2008; Gmeiner et al., 2013). Dopamine is most probably the strongest wake-promoting neuromodulator in fruit flies (reviewed by Birman, 2005). Hyperactive and sleepless *fumin* mutants carry a mutation in the dopamine transporter, which transports released dopamine back into the dopaminergic neurons (Kume et al., 2005). The *fumin* mutation results in a hypomorphic transporter, which leads to permanently high dopamine levels that continue to activate dopamine receptors on the postsynaptic neurons. Similar wake-promoting and sleep-reducing effects are observed when dopaminergic neurons are excited (Lima and Miesenböck, 2005; Wu et al., 2008, Shang et al., 2011; Liu et al., 2012; Ueno et al., 2012). Conversely, mutants deficient for tyrosine hydroxylase (TH), the rate-limiting enzyme for dopamine synthesis in the nervous system, have reduced dopamine levels and increased sleep throughout the day (Riemensperger et al., 2011).

In *D. melanogaster* the mushroom bodies (Joiner et al., 2006; Pitman et al., 2006; Yuan et al., 2006), the pars intercerebralis (Foltenyi et al., 2007; Crocker et al., 2010) and lateralis (Chen et al., 2016), the fan-shaped body of the central complex (Liu et al., 2012; Ueno et al., 2012; Pimentel et al., 2016; Donlea et al., 2018) have been identified as brain regions that regulate sleep. In addition, the Pigment-Dispersing Factor (PDF)-expressing large and small ventral Lateral Neurons (l-LN_v_s and s-LN_v_s), which belong to the circadian clock neurons have been identified as wake-promoting neurons within the flies circadian clock neuron network (Parisky et al., 2008; Sheeba et al., 2008a; Shang et al., 2008; Lebestky et al., 2009; Guo et al., 2016; Guo et al., 2018; Potdar and Sheeba, 2018; Liang et al., 2019).

The l-LN_v_s respond to both dopamine and octopamine through increases in cAMP, but the responses to dopamine are clearly stronger (Shang et al., 2011). Furthermore, the l-LN_v_s are directly light sensitive and promote arousal and activity in response to light, especially in the morning (Shang et al., 2008; Sheeba et al., 2008b; Fogle et al., 2011). Despite the strong responses of the l-LN_v_s to dopamine and their proposed role in controlling arousal, it is not known how dopamine-signaling to the l-LN_v_s increases wakefulness and inhibits sleep. Receptivity to dopamine in the s-LN_v_s has not been previously addressed. Here, we down-regulated the activating D1-like dopamine receptors Dop1R1 and Dop1R2 in the wake promoting l- and s-LN_v_s and examined the consequences on intracellular cAMP levels, resting membrane potential, and electrical firing rate in the electrophysiologically accessible l-LN_v_s. Moreover, we analyzed the behavioral consequences of Dop1R1/ Dop1R2 knock-down in the land s-LN_v_s on sleep and activity rhythms. As expected, we find that the knockdown of Dop1R1 reduces cAMP and electrophysiological responses to dopamine in the l-LN_v_s, confirming that dopamine signals via Dop1R1 receptors. Unexpectedly, we find that the down-regulation of the excitatory Dop1R1 receptor slightly decreases daytime sleep, suggesting that dopamine signaling via Dop1R1 to the LN_v_s usually promotes daytime sleep rather than wakefulness. Finally, we find that dopamine also likely signals to the s-LN_v_s via Dop1R2 receptors, and that the down-regulation of these receptors decreases night-sleep. Collectively, these results cast doubt on the currently held view of LN_v_s as dedicated wake-promoting neurons, and suggest a more complex regulation of sleep by these important clock neurons.

## Material and Methods

### Fly stocks

Flies were raised on *Drosophila* food (0.8 % agar, 2.2 % sugar-beet syrup, 8.0 % malt extract, 1.8 % yeast, 1.0 % soy flour, 8.0 % corn flour and 0.3 % hydroxybenzoic acid) at 25 °C under a 12 h:12 h light:dark (LD) cycle and transferred to 20 °C at an age of ~3 days.

To visualize TH-positive (dopaminergic) and the PDF-positive neurons we used *TH-Gal4* (Friggi-Grelin et al., 2002) to drive *UAS-10xmyrGFP* in dopaminergic neurons and stained with anti-GFP and anti-PDF. For visualizing presynapses of dopaminergic neurons and postsynapses of PDF neurons, we expressed the vesicle marker synaptotagmin::GFP (*UAS-sytI/II::GFP;* Bloomington) under control of *TH-Gal4* in dopaminergic neurons and a GFP labeled postsynaptic protein – the Down syndrome cell-adhesion molecule (*UAS-dscam::GFP*; Wang et al. 2004) – under control of *Pdf-Gal4* in PDF neurons. To visualize the spatial vicinity of dopaminergic and PDF fibers we used Split-GFP imaging (= GFP Reconstitution Across Synaptic Partners (GRASP); Feinberg et al., 2008): *yw;pdf-LexA/LexAop-GFP11;TH-Gal4/UAS-GFP1-10* flies were used to express the GFP11 fragment in the PDF-expressing LN_v_s and the GFP1-10 fragment in dopaminergic neurons, respectively. *yw;pdf-LexA/LexAop-GFP11;TM6B.Tb/UAS-GFP1-10* flies were used as controls.

In order to down-regulate the different dopamine receptors in all clock neurons or only in the PDF neurons (s-LN_v_s and l-LN_v_s), we used *Clk856-Gal4* (Gummadova et al., 2009) or *Pdf-Gal4* (Park et al., 2000), respectively to either express *UAS-Dop1R1_RNAi_* (no. 31765, Bloomington stock center), *UAS-Dop1R2_RNAi_* (no. 26018, Bloomington stock center) or *UAS-D2R_RNAi_* (no. 26001, Bloomington stock center) alone, or to simultaneously express *UAS-Dop1R1_RNAi_* and *UAS-Dop1R2_RNAi_*. The flies with the relevant Gal4 and UAS constructs (crossed with UAS-dicer2 flies) were taken as controls. In addition, we used an inducible Gal4 version, termed GeneSwitch (GS) (Osterwalder et al., 2001), under the control of the *Pdf* promotor (Depetris-Chauvin et al., 2011) to down-regulate Dop1R1 or Dop1R2 receptors in the PDF neurons only during adulthood of the flies. GS is a fusion between the Gal4 binding, the NFØb activation and the human progesterone receptor ligand-binding domains, which is expressed in the pattern dictated by the desired promoter but remains transcriptionally silent in the absence of RU486 (RU), an analog of progesterone. RU was mixed to the food of the adult flies in the Trikinetics monitors (see below). In all experiments *UAS-Dicer2* (no. 60012, Vienna *Drosophila* RNAi Center, Wien, Austria) was expressed additionally to enhance the effect. For simplicity we will call the experimental flies Clk856>Dop1Rx_RNAi_, Pdf>Dop1Rx_RNAi_ or PDF-GS>Dop1Rx_RNAi_, where the ‘x’ stands for the relevant dopamine receptor. Their sleep and activity profiles will always be depicted in red, while the relevant control flies are shown in black.

For imaging experiments the above described *Clk856-Gal4* or *Pdf-Gal4* line was used to express the ratiometric cAMP sensor *UAS-Epac1-camps* (Nikolaev et al., 2004), *UAS-dicer2* and the RNAi-constructs for different dopamine-receptors (see above).

### Immunostaining and microscopy

For immunostaining, whole-mount brains were fixed in 4 % paraformaldehyde (PFA) in phosphate buffered saline (PBS) for 2 hours at RT, followed by 4 washes in PBS containing 0.3 % TritonX-100 (PBT). They were blocked in 5 % normal goat serum (NGS) in PBT. Subsequently, the specimens were incubated in the primary antibody solution overnight at 4 °C. The primary antibody solution contained GFP antibody (raised in rabbit, Molecular Probes, A11122; dilution 1:1000) and PDF antibody (monoclonal mouse C7 antibody; Developmental Studies Hybridoma Bank at the University of Iowa; dilution 1:100). After rinsing in PBT, fluorescence conjugated secondary antibodies (Alexa-Fluor^®^ Dyes, Molecular Probes, Carlsbad, CA) were applied overnight at 4 °C. The stained brains were finally embedded in Vectashield and scanned with a Confocal Microscope (Leica TCS SPE, Wetzlar, Germany).

### Ex vivo live-cAMP imaging

Flies were well entrained to a LD 12:12 cycle and imaging always took place during the light phase of the LD cycle (between ZT2 and ZT8). For imaging, flies were anesthetized on ice and brains were dissected in cold hemolymph-like saline (HL3; Stewart et al., 1994) and mounted at the bottom of a plastic petri dish in HL3. Brains were allowed to recover from dissection for at least 10 min prior to imaging. An epifluorescent imaging setup (VisiChrome High Speed Polychromator System, ZEISS Axioskop2 FS plus, Visitron Systems GmbH) with a 40x dipping objective (ZEISS 40x/1,0 DIC VIS-IR) was used for all imaging experiments. Neurons were localized using GFP-optics and were identified according to their position in the brain. Regions of interest were defined on single cell bodies in the Visiview Software (version 2.1.1, Visitron Systems GmbH). Time-lapse frames were acquired with 0.2 Hz for 12 min, exciting the CFP fluorophore of the ratiometric cAMP sensor with light of 405 nm. Emissions of CFP and YFP were detected separately by a CCD-camera (Photometrics, CoolSNAP HQ, Visitron Systems GmbH) with a beam splitter. After measuring baseline CFP and YFP levels for ~100 s, pharmacological treatments were bath applied drop-wise using a pipette. HL3 application served as negative control and 10 μM NKH^477^ (an activator of all adenylate cyclases) as positive control. Dopamine and SKF^38393^ (a DopR1 agonist) were diluted in HL3 and were applied in an end concentration of 1 mM and 0.1 mM, respectively. For Tetrodotoxin (TTX)-treatments, brains were incubated in 2 μM TTX in HL3 for 20 min prior to imaging and dopamine was diluted in 2 μM TTX in HL3 for the application. Inverse Fluorescence Resonance Energy Transfer (iFRET) was calculated according to the following equation: iFRET=CFP/(YFP-CFP*0.357) (Shafer et al., 2008). Thereby, CFP and YFP are background corrected raw fluorescence data and 0.357 was determined as the fraction of CFP spillover into the YFP channel in our imaging setup, which had to be subtracted from YFP fluorescence. Finally, iFRET traces of individual neurons were normalized to base line levels and were averaged for each treatment. For quantification and statistical comparison of response amplitudes of each treatment or genotype, maximum iFRET changes were determined for individual neurons.

### Ex vivo patch-clamp electrophysiology

Three to nine days-old female flies were anesthetized with a brief incubation of the vial on ice, brain dissection was performed in external recording solution which consisted of (in mM): 101 NaCl, 3 KCl, 1 CaCl_2_, 4 MgCl_2_, 1.25 NaH_2_PO_4_, 5 glucose, and 20.7 NaHCO_3_, pH 7.2, with an osmolarity of 250 mmol/kg (based on saline solution used by Cao and Nitabach, 2008). After removal of the proboscis, air sacks and head cuticle, the brain was routinely glued ventral side up to a sylgard-coated coverslip using a few microliters of tissue adhesive 3 M Vetbond. The time from anesthesia to the establishment of the recordings was approximately 20 minutes spent as following: l-LN_v_s were visualized by red fluorescence in *Pdf-RFP* flies (which express a red fluorophore under the Pdf promoter, Ruben et al., 2012) using an Olympus BX51WI upright microscope with 60X water-immersion lens and ThorLabs LEDD1B and TK-LED (TOLKET S.R.L, Argentina) illumination systems. Once the fluorescent cells were identified, cells were visualized under IR-DIC using a DMK23UP1300 Imaging Source camera and IC Capture 2.4 software. l-LN_v_s were distinguished from s-LN_v_s by their size and anatomical position. To allow the access of the recording electrode, the superficial glia directly adjacent to l-LN_v_s somas was locally digested with protease XIV solution (10 mg/ml, SIGMA-ALDRICH P5147) dissolved in external recording solution. This was achieved using a large opened tip (approximately 20 μm) glass capillary (pulled from glass of the type FG-GBF150-110-7.5, Sutter Instrument, US) and gentle massage of the superficial glia with mouth suction to render the underling cell bodies accessible for the recording electrode with minimum disruption of the neuronal circuits. After this procedure, protease solution was quickly washed by perfusion of external solution. Recordings were performed using thick-walled borosilicate glass pipettes (FG-GBF150-86-7.5, Sutter Instrument, US) pulled to 7-8 MΩ using a horizontal puller P-97 (Sutter Instrument, US) and fire polished to 9-12 MΩ. Recordings were made using a Multiclamp 700B amplifier controlled by pClamp 10.4 software via an Axon Digidata 1515 analog-to-digital converter (Molecular Devices, US). Recording pipettes were filled with internal solution containing (in mM): 102 potassium gluconate, 17 NaCl, 0.085 CaCl_2_, 0.94 EGTA and 8.5 HEPES, pH 7.2 with an osmolarity of 235 mmol/kg (based on the solution employed by Cao and Nitabach 2008). Gigaohm seals were accomplished using minimal suction followed by break-in into whole-cell configuration using gentle suction in voltage-clamp mode with a holding voltage of −60 mV. Gain of the amplifier was set to 1 during recordings and a 10 kHz lowpass filter was applied throughout. Spontaneous firing was recorded in current clamp (I=0) mode. Analysis of traces was carried out using Clampfit 10.4 software. For action potential firing rate calculation the event detection tool of Clampfit 10.4 was used. Perfusion of external saline in the recording chamber was achieved using a peristaltic pump (Ismatec ISM831). After 3 min of recording basal conditions, 10 ml of Dopamine (1 mM) prepared in external saline were perfused, this lasted approximately 3 minutes. Dopamine was then washed out with external saline perfusion during 10 minutes. For basal condition, the number of action potentials on the last minute before Dopamine application was counted. For Dopamine condition, the number of action potentials was counted on the last minute of Dopamine perfusion. For wash out condition, the number of action potentials was counted on the last minute of the recording. In all cases, the firing rate in Hz was calculated by dividing the number of action potentials over 60 seconds. The membrane potential was assessed during the same periods for each condition. All recordings were performed during the time-range of ZT6 to ZT9.

### Recording of sleep and activity

Locomotor activity of male 3-7 days old flies was recorded as described previously (Hermann-Luibl et al., 2014) using *Drosophila Activity* Monitors by TriKinetics. The fly tubes were fixed by a Plexiglas frame in such a way that the infrared beam crossed each fly tube at a distance of ~3 mm from the food. The food consisted of 4 % sugar in agar. For the gene-switch experiments, RU486 (mifepristone, Sigma) was dissolved in 80 % ethanol and mixed with the food to a final concentration of 200 mg/ml. In the controls the same amount of ethanol (vehicle) was added to the food. Flies were monitored for 9 days in 12 h:12 h light-dark cycles 12:12 (LD 12:12) with a light intensity of 100 lux at 20 °C and then released into constant darkness (DD). Recording days 3-7 in LD were used for sleep and activity analysis.

Sleep analysis was performed with a custom-made Excel Macro (provided by T. Yoshii; Gmeiner et al., 2013; Hermann-Luibl et al., 2014). Sleep was defined as the occurrence of 5 consecutive recording minutes without interruption of the infrared-beam within the TriKinetics monitor. For average daily sleep profiles, sleep was calculated in 1-hour-bins and averaged over the 5 selected days for each single fly and genotype. Furthermore, the total amount of sleep was averaged over the 5 days, as well as the amount of sleep during the light phase and the dark phase and the average sleep bout duration. Every experiment was repeated at least twice and at a minimum 30 flies of each genotype were used for the analysis.

The same 5 days of recordings used for sleep evaluation were also analyzed for fly activity. Daily average activity profiles were calculated for each fly as described in Schlichting and Helfrich-Förster (2015). From these, the total activity (number of infrared-beam crosses) of every fly during the entire day, the dark-phase and the light-phase were calculated and plotted for each genotype. An activity index (the average of beam crosses per active minute) was also calculated but not shown, since it correlated with the total activity. The free-running period of each fly was determined from the recordings in DD to judge whether downregulating the dopamine receptors changed the speed of the circadian clock.

### Statistics

Statistical analyses of sleep and activity data were performed using the R environment (v3.5.3). Data were tested for normal distribution with a Shapiro-Wilk normality test (p>0.05). The three data groups “whole day”, “day” and “night” were tested separately. If any group wasn’t normally distributed the whole dataset was handled as not normally distributed. In this case the Mann–Whitney *U* test was used. A T-test was used for normally distributed data in case of variance homogeneity (Levene’s test, p>0.05). Period length was tested for statistically significant influences of dopamine receptor RNAi and RU treatment by a two-way ANOVA followed by a post-hoc test with Bonferroni correction. Statistical tests on live imaging data were also done with the R environment. We compared the Epac1-camps inverse FRET ratio between vehicle and test compounds and used the Wilcoxon signed rank test with Bonferroni correction for multiple comparisons of maximum changes. Exceptions are stated in the figure legends. Electrophysiological data (membrane potential and firing rate) was analyzed with Kruskal-Wallis non-parametric test, the alpha parameter was 0.05 and the post hoc test used the Fisher’s least significant difference criterion. Bonferroni correction was applied as the adjustment method.

## Results

### Dopaminergic neurons are presynaptic to the ventrolateral clock neurons (l-LN_v_s and s-LN_v_s) that arborize in the accessory medulla

Both, the s-LN_v_s and l-LN_v_s express the neuropeptide PDF and send dendrites into the accessory medulla (AME) – the insect clock center (Helfrich-Förster, 1995; Helfrich-Förster et al., 2007). These neurons are thought to be wake-promoting: their activity coincides with the morning peak of wakefulness (Liang et al., 2019), and their optogenetic excitation, along with other lateral neuron types, reduces sleep (Guo et al., 2018). The s-LN_v_s project into the dorsolateral brain and are there connected to other clock neurons and several neurons downstream of the clock that control activity and sleep (reviewed in King and Sehgal, 2020). The l-LN_v_s are conspicuous clock neurons with wide arborizations in the ipsilateral and contralateral optic lobe and connections between the brain hemispheres (Helfrich-Förster et al., 2007). In the AME, their neurites overlap with those of dopaminergic neurons (Hamasaka and Nässel, 2006; Shang et al., 2011). Microarray studies show that they express genes encoding the excitatory dopamine receptors Dop1R1, Dop1R2, and DopEcR) and the inhibitory dopamine D2R, in addition to the excitatory octopamine receptors OAMB and OA2 (Kula-Eversole et al., 2010; Shang et al., 2011). The AME of *Drosophila* can be subdivided into two parts: a central part and a ventral elongation (Fig. 1). Whereas the central part is innervated by several clock neurons including the PDF-positive small ventrolateral neurons (s-LN_v_s), the ventral elongation only receives fibers from the l-LN_v_s (Helfrich-Förster et al., 2007; Schubert et al., 2018). Previous studies already suggested that the PDF-fibers in the ventral elongation of the AME are predominantly postsynaptic (of dendritic nature) (Helfrich-Förster et al., 2007) and in close vicinity to dopaminergic fibers (Shang et al., 2011; Fig. 1a), but whether the dopaminergic fibers were of presynaptic nature was unclear. By expressing the vesicle marker *Synaptotagmin (SytI/II)::GFP* in the *TH-Gal4*-positive (dopaminergic) neurons and the postsynaptic marker *Dscam::GFP* in the *Pdf-Gal4*-positive neurons we show here that this is indeed the case (Fig. 1). Prominent SytI/II::GFP staining was present in *TH-Gal4*-positive fibers that are aligned along the ventral elongation (Fig. 1c) and Dscam::GFP was strongly localized in the PDF fibers of the entire ventral elongation of the AME (Fig. 1d). Using GRASP imaging, we confirmed previous results that PDF-and *TH-Gal4*-positive fibers have contact in the central part of the AME and its ventral elongation (Shang et al., 2011): reconstituted GFP signals were present in both parts of the AME (Fig. 1b), whereas no reconstituted GFP signals were detected in control flies. In summary, we show here that the dopaminergic neurons are presynaptic to the l-LN_v_s and s-LN_v_s.

**Figure 1.**
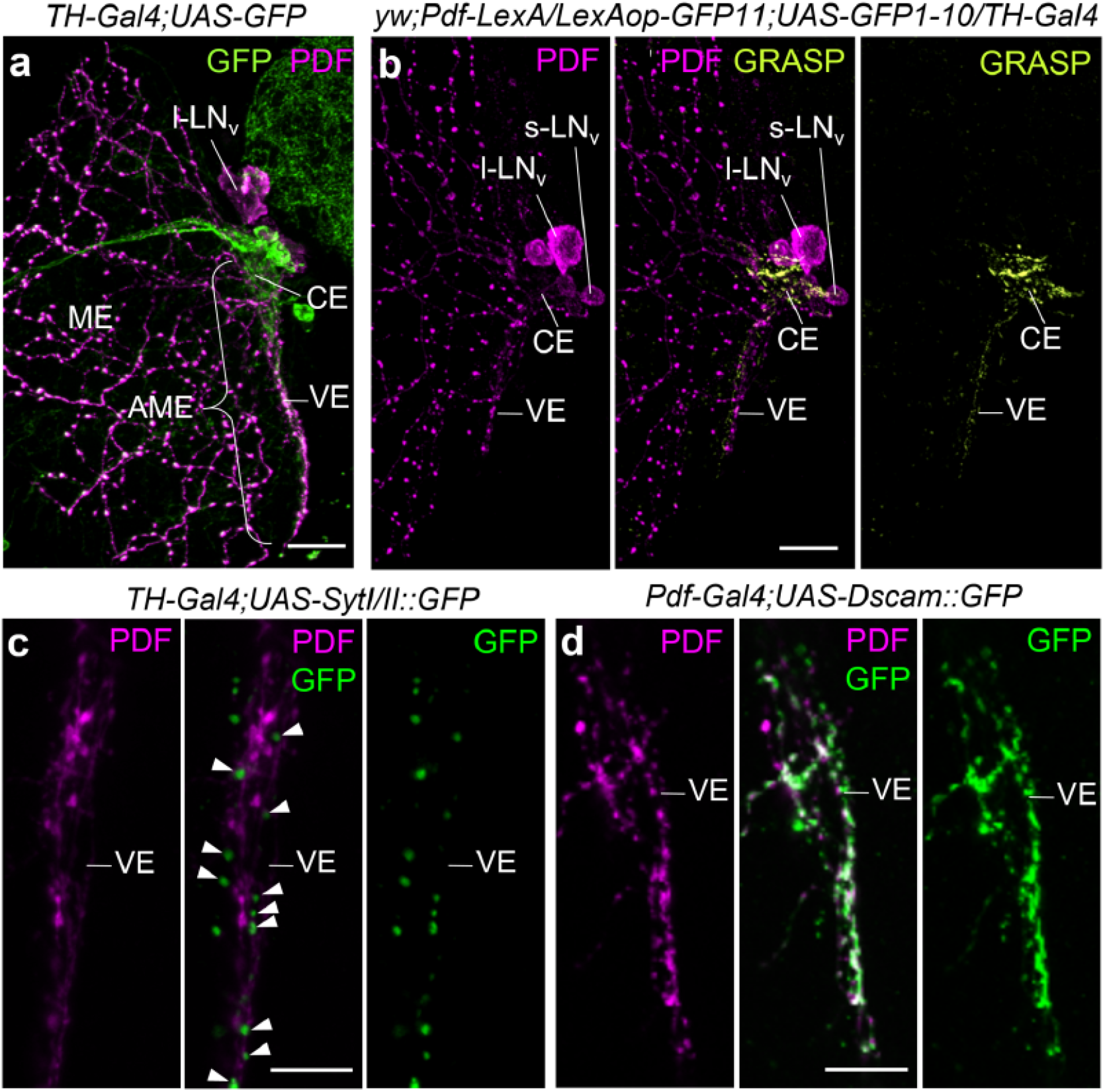
Staining of whole-mount brains showing the spatial vicinity of dopaminergic neurites (visualized with *TH-Gal4*) and neurites from the PDF-positive LN_v_s in the accessory medulla of one hemisphere. All pictures are overlays of 2 μm thick confocal stacks. (**a**) Medulla (ME) and accessory medulla (AME) labeled with anti-PDF (magenta) and anti-GFP (*TH-Gal4;UAS-10xmyrGFP,* green) (overlay of 10 confocal stacks). *TH-Gal4* and PDF overlap in the central part (CE) and ventral elongation (VE) of the AME. l-LN_v_s, PDF-positive large ventrolateral neurons; s-LN_v_s, PDF-positive small ventrolateral neurons. (**b**) GFP Reconstitution Across Synaptic Partners (GRASP) between *Pdf-Gal4* neurons and *TH-Gal4* neurons. GRASP signals are found in the CE and VE of the AME (overlay of 6 confocal sections). (**c**) Expression of the presynaptic marker Synaptotagmin::GFP (SytI/II::GFP) in the *TH-Gal4* neurons (GFP; green) and co-staining against PDF (magenta) (overlay of 3 confocal stacks). GFP-positive vesicles (arrowheads) are present along the PDF-positive fibers in the VE. (**d**) Expression of the postsynaptic marker Dscam::GFP (green) in the *Pdf-Gal4*-positive l-LN_v_s and co-staining with anti-PDF (magenta) (overlay of 3 confocal stacks). The PDF-positive fibers in the VE of the AME are predominantly dendritic. Scale bars = 20 μm in **a** and **b**, and 10 μm in **c** and **d**.

### Dopamine signals to different clock neurons

It was shown previously that dopamine application to isolated brains elevates cAMP levels in the l-LN_v_s (Shang et al., 2011). We confirmed this result and extended it to the other clock neurons that have arborizations in the central part of the AME, i.e. the s-LN_v_s, the dorsolateral neurons (LN_d_s) and the anterior dorsal neurons 1 (DN_1a_s) (Helfrich-Förster et al., 2007; Schubert et al., 2018). The l-LN_v_s showed the strongest responses to dopamine, which were even higher after blocking synaptic transmission by TTX, suggesting that inhibitory signals from other interneurons usually reduce the cAMP response to dopamine (Fig. 2a). Significant responses to dopamine that persisted under TTX were also present in the LN_d_s (Fig. 2b) and the DN_1_s (Fig. 2c). The s-LN_v_s also exhibited significantly increased cAMP levels after dopamine application; but these cells are hard to image, because they are very small and often located underneath the l-LN_v_s, so that their responses cannot be unequivocally separated from those of the l-LN_v_s. Therefore, we could only image a few of them without application of TTX (Fig. 3).

**Figure 2.**
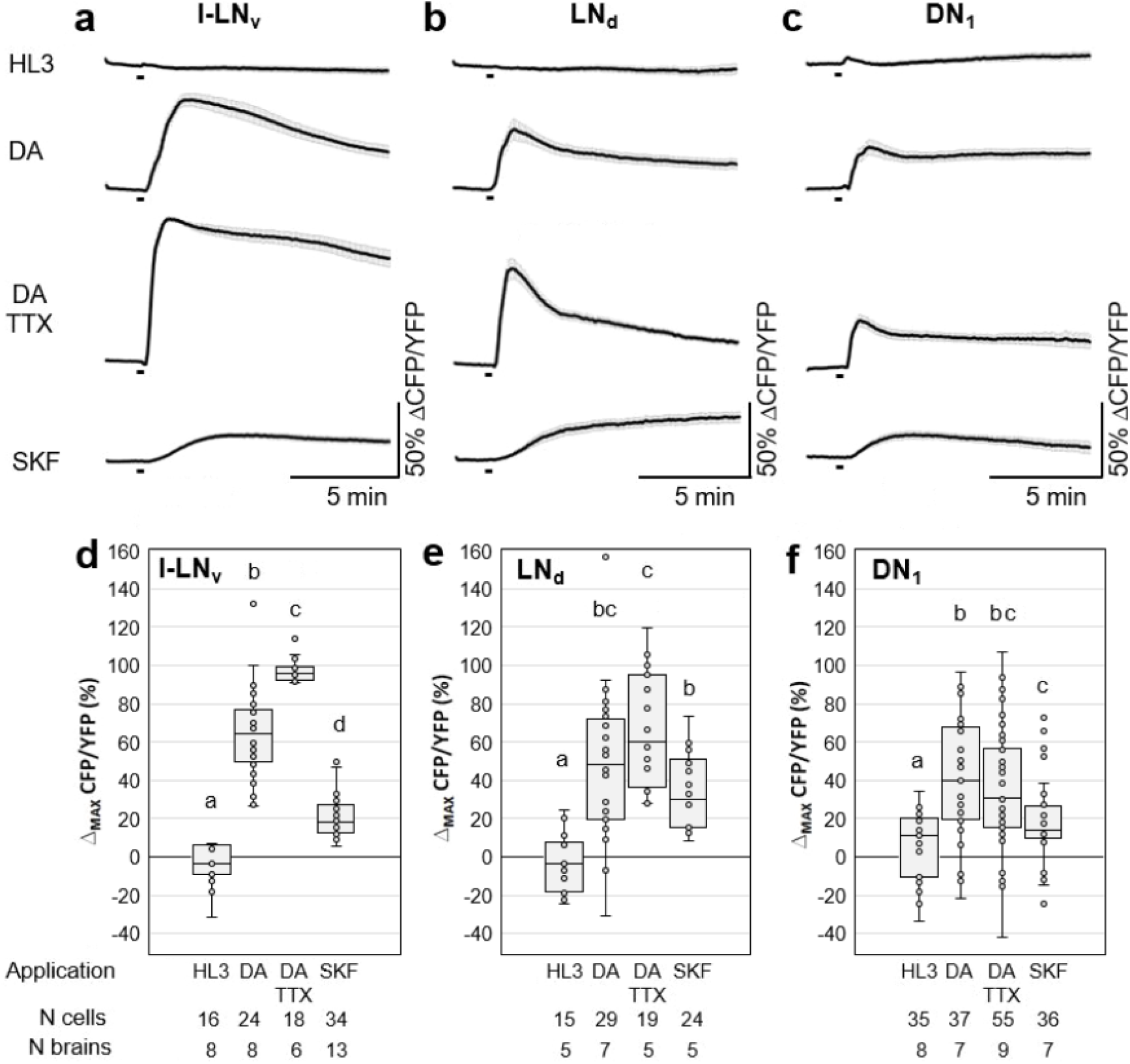
*Ex vivo* live-cAMP imaging on *Drosophila* clock neurons. (**a-c**) Mean inverse FRET traces of l-LN_v_, LN_d_ and DN_1_ clock neurons of *clk856> Epac1* flies. Error bars (grey) represent SEM and short black bars indicate application of the different solutions: HL3 = buffer (negative control), DA (= 1 mM dopamine), DA+TTX (= 1 mM DA + 2 μM Tetrodotoxin) and SKF^38393^ (= 0.1 mM Dop1R1-agonist), respectively. (**d-f**) Quantification of maximum inverse FRET changes for each single neuron (dots in Box Plots) of each treatment. Black horizontal lines in the Box Plots represent the median, different letters indicate significant differences. Cells of all three neuronal clusters respond with robust and significant increases in cAMP levels upon application of DA and DA+TTX compared to negative controls, indicating a direct neuronal connection between dopaminergic neurons and clock neurons. Application of the Dop1R1-agonist SKF also significantly increased cAMP levels in all three clusters of clock neurons (**f**).

**Figure 3.**
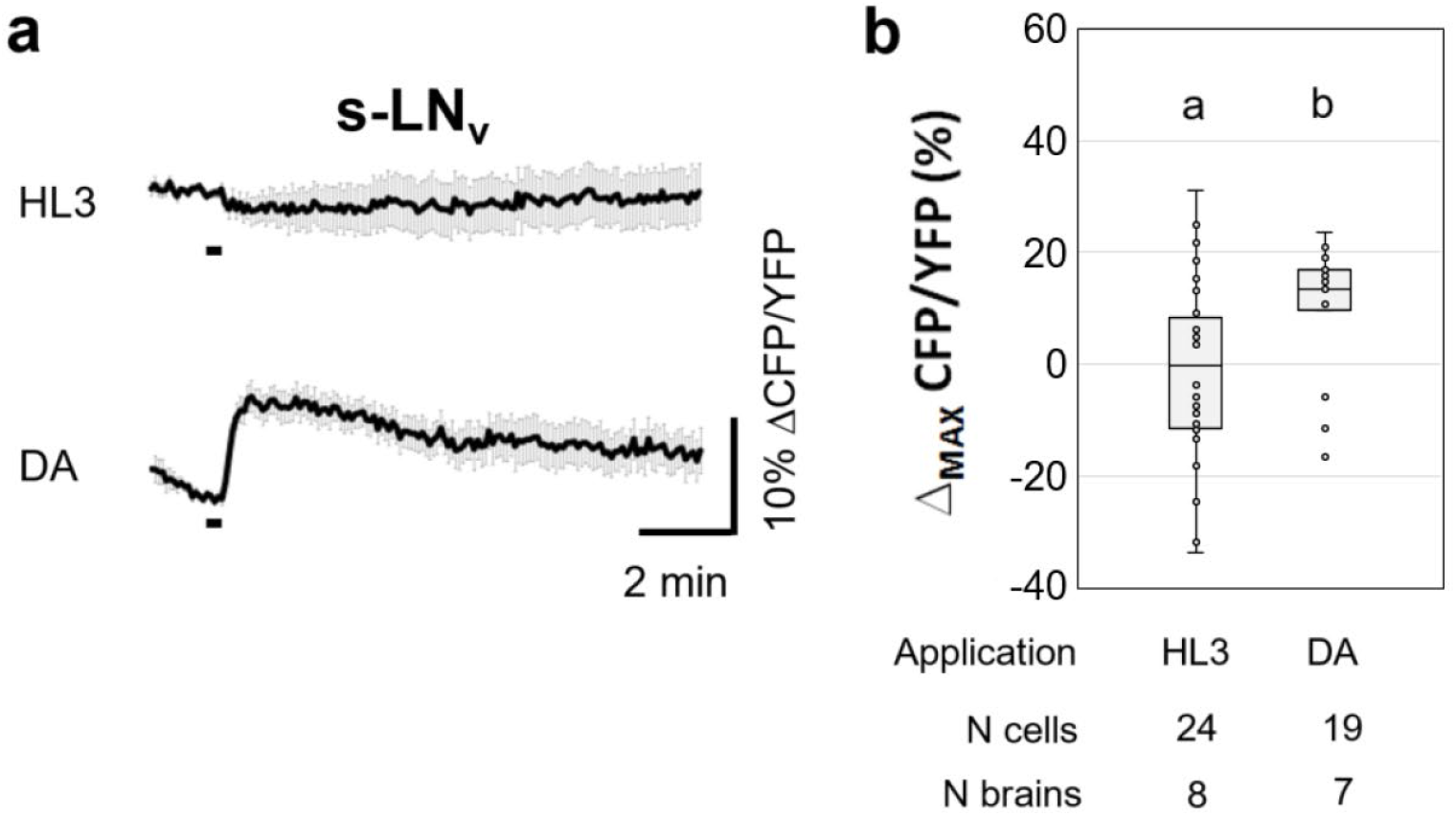
Ex vivo live-cAMP imaging on Drosophila s-LNv neurons. (**a**) Mean inverse FRET traces of s-LNv clock neurons of clk856>Epac flies. Error bars (grey) represent SEM and short black bars indicate application of negative control (HL3) or 1 mM dopamine (DA). (**b**) Quantification of maximum inverse FRET changes for each single neuron (dots in Box Plots) of each treatment. Black horizontal lines in the Box Plots represent the median, different letters indicate significant differences. s-LN_v_s significantly responded to DA with an increase in cAMP. In this case the Mann–Whitney U test was used for pairwise comparison of maximum changes.

Next, we tested whether these cAMP responses were mediated by Dop1R1 or Dop1R2 receptors. Knockdown of Dop1R1 by RNAi in all clock neurons, reduced cAMP responses in the l-LN_v_s (Fig. 4a, d), the DN_1_s (Fig. 4c, f) and the LN_d_s (Fig. 4b, e), whereas the down-regulation of Dop1R2 appeared to reduce cAMP levels in all neuron clusters slightly but not significantly (Fig. 4a-c). Notably, the cAMP signals in the LN_d_s were quite variable when Dop1R1 or Dop1R2 were down-regulated; some neurons still responded to dopamine, while others did not (Fig. 4e). The same applies for the DN_1_s knockdown of Dop1R1; half of the cells responded, the other half did not (Fig. 4f). However, with knockdown of Dop1R2, only two of the measured 22 DN_1_ cells did not respond to dopamine (Fig. 4f). Altogether, this suggests that some LN_d_s and DN_1_s express Dop1R1 and others Dop1R2. Consistent with this hypothesis the simultaneous downregulation of Dop1R1 and Dop1R2 abolished the responses to dopamine in all evaluated neurons (Fig. 4). Down-regulation of the inhibitory dopamine receptor D2R, slightly increased the responses to dopamine in the l-LN_v_s (Fig. 4a, d) and the LN_d_s (Fig. 4b, e); but in contrast to a previous study (Shang et al., 2011) this increase was not significant. To make sure that the neurons were able to increase their cAMP levels in our setup, we measured cAMP levels in responses to NKH^477^, an adenylyl cyclase activator, and found that they all responded (Fig. 5).

**Figure 4.**
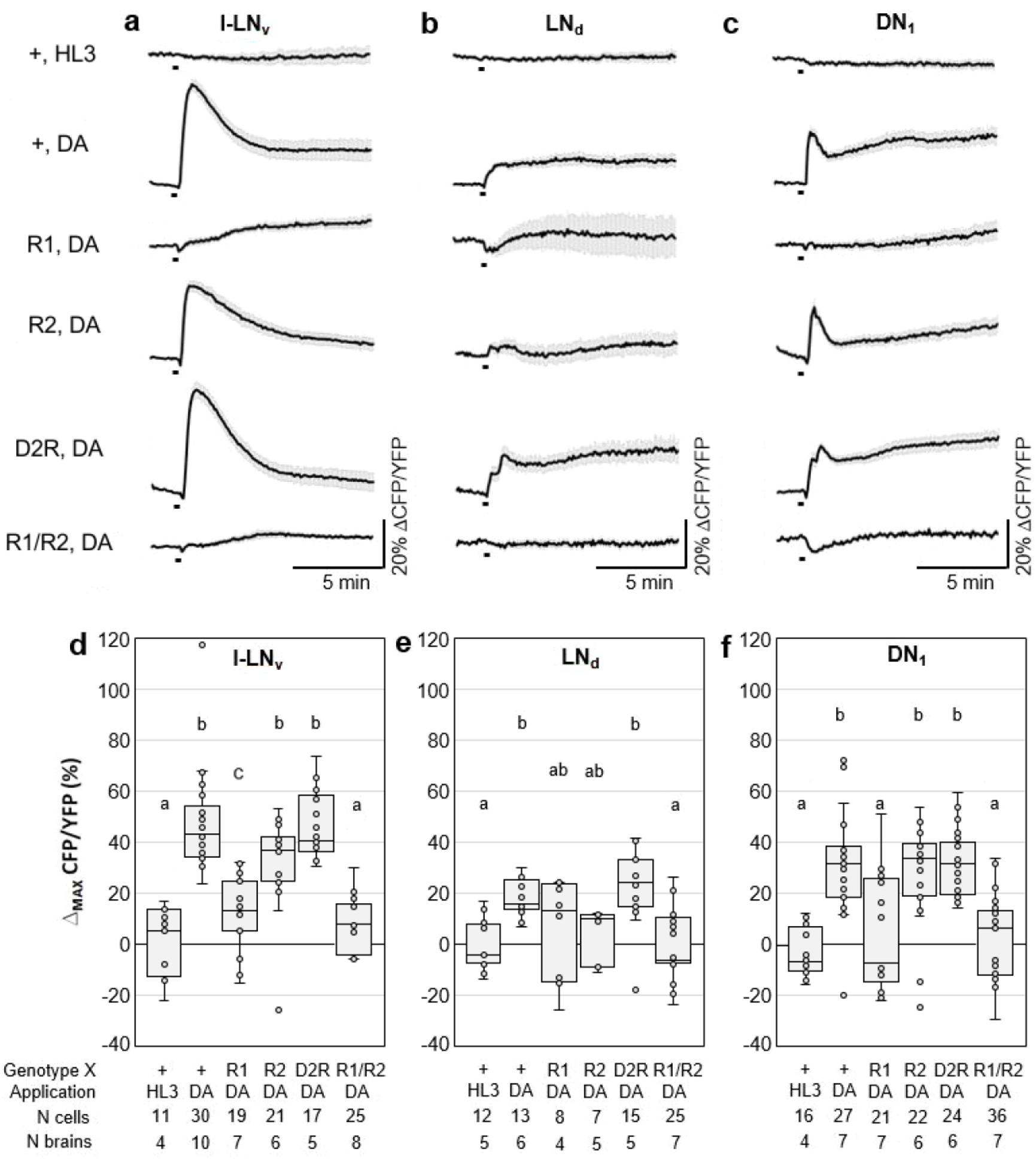
*Ex vivo* live-cAMP imaging on *Drosophila* clock neurons expressing RNAi-constructs against different Dopamine receptors. (**a-c**) Mean inverse FRET traces of l-LN_v_, LN_d_ and DN_1_ clock neurons of *clk856>dicer2, Epac1, X_RNAi_* flies. The X stands for ‘wildtype’ (+) or the relevant dopamine receptor RNAi lines: R1 = *Dop1R1_RNAi_*, R2 = *Dop1R2_RNAi_*, D2R = *D2R_RNAi_* and R1/R2 = *Dop1R1_RNAi_/Dop1R2_RNAi_*. Error bars (grey) represent SEM and short black bars indicate application of negative control (HL3) or 1 mM dopamine (DA). (**d-f**) Quantification of maximum inverse FRET changes for each single neuron (dots in Box Plots) of each treatment. Black horizontal lines in the Box Plots represent the median, different letters indicate significant differences. DN_1_ neurons responded significantly to application of DA, except when Dop1R1 or Dop1R1/R2 were knocked down. l-LN_v_ neurons lacked the responses to dopamine when both dopamine receptors (Dop1R1/R2) were knocked down Responses of the LN_d_ were not different from negative controls when either Dop1R1 or Dop1R2 or both (Dop1R1/R2) were knocked down.

**Figure 5.**
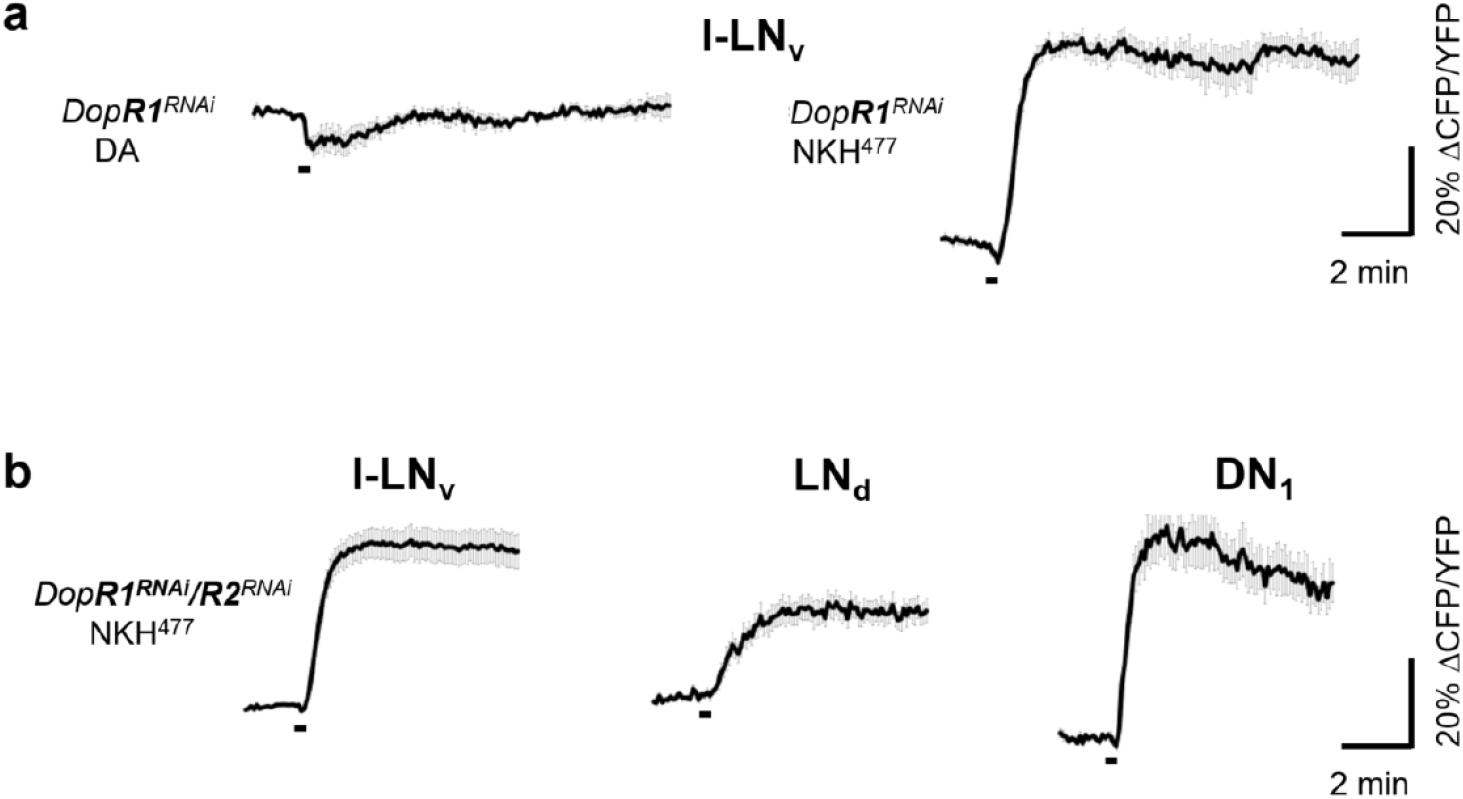
*Ex vivo* live-cAMP imaging on *Drosophila* clock neurons expressing RNAi-constructs against different dopamine receptors (*clk856>dicer2,Epac1;Dop1Rx_RNAi_* flies). (**a**) Mean inverse FRET traces of l-LN_v_ clock neurons with down-regulated Dop1R1 (*Dop1R1_RNAi_*). The same set of neurons (5 neurons from 2 brains) was first subject to 1 mM dopamine (DA) application showing no response and afterwards to application of 10 μM of the adenylate-cyclase activator NKH^477^, which evoked an increase in cAMP. (**b**) Mean inverse FRET traces of the same l-LN_v_, LN_d_ and DN_1_ clock neurons shown in Fig. 4a, b, c (bottom) expressing *DopR1_RNAi_/DopR2_RNAi_* after application of NKH^477^. Error bars (grey) represent SEM and short black bars indicate application of negative control (HL3) or 1 mM dopamine (DA).

In summary, our results show that the responses to dopamine are predominantly mediated by Dop1R1 receptors in the l-LN_v_s and DN_1_s and by Dop1R1 and Dop1R2 receptors in the LN_d_s. As described above, we could not identify the relevant Dop1R1 receptors of the s-LN_v_s, because these cells were hidden by the l-LN_v_s or just located too close to them, which prevented a successful imaging in all the preparations with down-regulated Dop1R receptors.

### Effects of Dop1R1 and Dop1R2 down-regulation in the clock neurons on sleep

To study the consequences of reduced dopamine signaling in the LN_v_ clock neurons on sleep, we first down-regulated the activating Dop1R1 and Dop1R2 receptors in all clock neurons (using *Clk856-Gal4*). We did not see any significant changes in sleep pattern (Fig. 6a), total sleep, or sleep during day and night, nor on sleep bout duration (Fig. 6b) with down-regulation of each of the receptors alone or down-regulation of both receptors simultaneously. However, the activity level during the day was significantly reduced by down-regulation of each of the two dopamine receptors alone or in combination (Fig. 6c, d). Furthermore, in the case of Dop1R2 down-regulation, activity during the night was significantly increased (Fig. 6d). The free-running period in constant darkness did not change when dopamine receptors were knocked down, only the power of the rhythm was decreased slightly by knockdown of both dopamine receptors simultaneously (Table 1).

**Figure 6.**
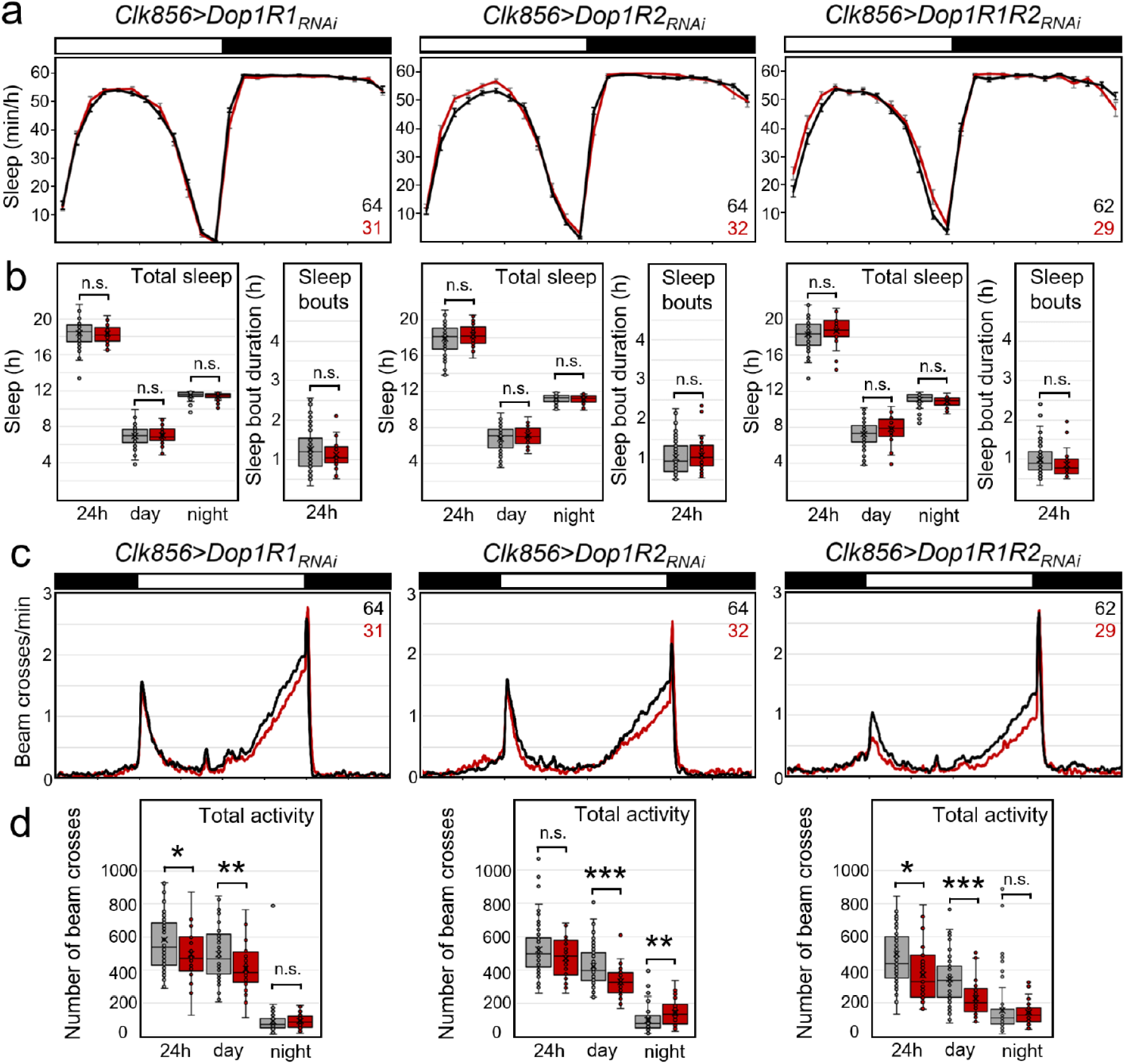
Sleep and activity in clock neuron specific dopamine-receptor knockdown flies. Dop1R1, Dop1R2 or both were knocked down using *clk856-Gal4*. (**a**) Average daily sleep profiles of experimental flies (red, *clk856>Dop1R_RNAi_*) and respective *Gal4* and *UAS* controls (pooled in black; controls were not significantly different from each other and were pooled in a single). (**b**) Box Plots of sleep parameters (total sleep in hours during the entire 24 h period, during the day and the night; same color code as in **a**). The median, upper and lower quartiles as well as upper and lower extremes plus the single data points are plotted. No significant differences were observed between experimental flies and controls in any of the three cases. (**c**) Average activity profiles of the same flies that are depicted in **a**. The flies with down-regulated dopamine receptors were always less active during day as compared to the controls. (**d**) Box Plots of total activity during the entire 24 h period, during the day and the night. Significant differences are indicated by asterisks (* p<0.05; ** p<0.01; *** p<0.001). The numbers of tested flies are indicated in (**a**) and (**c**).

**Table 1.** Rhythmic parameters of the free-running rhythms under constant darkness (DD)

| | rhythmic | period (± SD) | relative power (± SD) |
| --- | --- | --- | --- |
| *UAS-dicer2;clk856-Gal4;UAS-Dop1R1<sub>RNAi</sub>* | 94 % | 23.82 ± 0.41 | 3950.63 ± 1145.38 |
| *UAS-dicer2;clk856-Gal4;UAS-Dop1R2<sub>RNAi</sub>* | 100 % | 23.93 ± 0.40 | 4637.78 ± 1399.28 |
| *UAS-dicer2;clk856-Gal4;UAS-Dop1R1R2<sub>RNAi</sub>* | 84 % | 23.83 ± 0.44 | 2687.00 ± 772.08* |
| *UAS-dicer2;clk856-Gal4* | 97 % | 24.03 ± 0.46 | 3709.97 ± 1146.97 |
| *UAS-dicer2;Pdf-Gal4;UAS-Dop1R1<sub>RNAi</sub>* | 97 % | 23.93 ± 0.33 | 6496.61 ± 1652.04 |
| *UAS-dicer2;Pdf-Gal4;UAS-Dop1R2<sub>RNAi</sub>* | 97 % | 24.09 ± 0.41 | 5315.03 ± 2031.54 |
| *UAS-dicer2;Pdf-Gal4;UAS-Dop1R1R2<sub>RNAi</sub>* | 97 % | 24.00 ± 0.31 | 3752.74 ± 1485.88 |
| *UAS-dicer2;Pdf-Gal4* | 100 % | 24.31 ± 0.34 | 6966.94 ± 1699.23 |
| *UAS-dicer2;;UAS-Dop1R1<sub>RNAi</sub>* | 100 % | 23.70 ± 0.42 | 4812.78 ± 1702.03 |
| *UAS-dicer2;;UAS-Dop1R2<sub>RNAi</sub>* | 100 % | 23.75 ± 0.26 | 4198.81 ± 1351.07 |
| *UAS-dicer2;;UAS-Dop1R1R2<sub>RNAi</sub>* | 100 % | 23.66 ± 0.34 | 4135.06 ± 1298.63 |
| *UAS-dicer2;Pdf-GS;UAS-Dop1R1<sub>RNAi</sub> + Eth* | 100 % | 23.74 ± 0.35 | 2833.94 ± 720.94 |
| *UAS-dicer2;Pdf-GS;UAS-Dop1R1<sub>RNAi</sub> + RU* | 100 % | 24.18 ± 0.54** | 2668.28 ± 365.25 |
| *UAS-dicer2;Pdf-GS;UAS-Dop1R2<sub>RNAi</sub> + Eth* | 94 % | 23.84 ± 0.34 | 3886.97 ± 978.24 |
| *UAS-dicer2;Pdf-GS;UAS-Dop1R2<sub>RNAi</sub> + RU* | 97 % | 24.87 ± 0.52** | 2865.97 ± 461.19 |
| *UAS-dicer2;Pdf-GS + Eth* | 100 % | 23.76 ± 0.24 | 4120.34 ± 715.72 |
| *UAS-dicer2;Pdf-GS + RU* | 100 % | 24.46 ± 0.42** | 4332.97 ± 894.48 |
| *UAS-dicer2;;UAS-Dop1R1<sub>RNAi</sub> + Eth* | 100 % | 23.84 ± 0.34 | 3686.97 ± 978.24 |
| *UAS-dicer2;;UAS-Dop1R1<sub>RNAi</sub> + RU* | 100 % | 23.89 ± 0.25 | 3608.44 ± 830.90 |
| *UAS-dicer2;;UAS-Dop1R2<sub>RNAi</sub> + Eth* | 100 % | 23.81 ± 0.30 | 3433.91 ± 859.93 |
| *UAS-dicer2;;UAS-Dop1R2<sub>RNAi</sub> + RU* | 100 % | 23.81 ± 0.36 | 3526.28 ± 582.67 |

\* significant differences ( $p < 0.05$ ) in power between flies with down-regulated dopamine receptors in all clock neurons in comparison to the relevant controls
\*\* highly significant differences ( $p < 0.01$ ) after RU application in the *Pdf*-GeneSwitch (GS) experiments

Since among all clock neurons the s-LN_v_s and l-LN_v_s have been the ones with the most prominent role in sleep and arousal regulation, we decided to repeat Dop1R1 and Dop1R2 receptor down-regulation more specifically using the *Pdf-Gal4* driver. The l- and s-LN_v_s collectively produce the first daily peak of wakefulness (Renn et al., 1999; Grima et al., 2004; Stoleru et al., 2004; Rieger et al., 2006; Potdar and Sheeba, 2018; Liang et al., 2019) and the l-LN_v_s mediate light driven arousal (Parisky et al., 2008; Shang et al., 2008; Sheeba et al., 2008a; Lebestky et al., 2009). We repeated Dop1R1 and Dop1R2 receptor down-regulation in these neurons using the *Pdf-Gal4* driver. Once again, the general sleep pattern was not affected by the down-regulation (Fig. 7a), but total sleep and mean sleep bout duration were significantly reduced after all manipulations (down-regulation of Dop1R1 or Dop1R2 and simultaneous down-regulation of both receptors) (Fig. 7b). Closer inspection revealed that Dop1R1 downregulation reduced sleep significantly during the day, whereas Dop1R2 down-regulation reduced sleep significantly both during the day and night, as did the down-regulation of both receptors simultaneously. The effects of dopamine receptor down-regulation on activity levels were mixed. We did not observe any effects on daytime activity, but nighttime activity was slightly but significantly increased by Dop1R2 receptor knockdown and knockdown of both receptors (Fig. 7c, d). We did not observe any effects on the period or the power of the free-running rhythms in DD (Table 1). In summary, these results suggest that reduction in dopamine signaling in the LN_v_s has no effect on the speed of the clock. However, dopamine signaling unexpectedly appears to increase sleep via Dop1R1 receptors during the day and via Dop1R2 receptors during the day and the night. These results should be treated with caution because they were achieved by constitutive knockdown of dopamine receptors, which may cause developmental effects.

**Figure 7.**
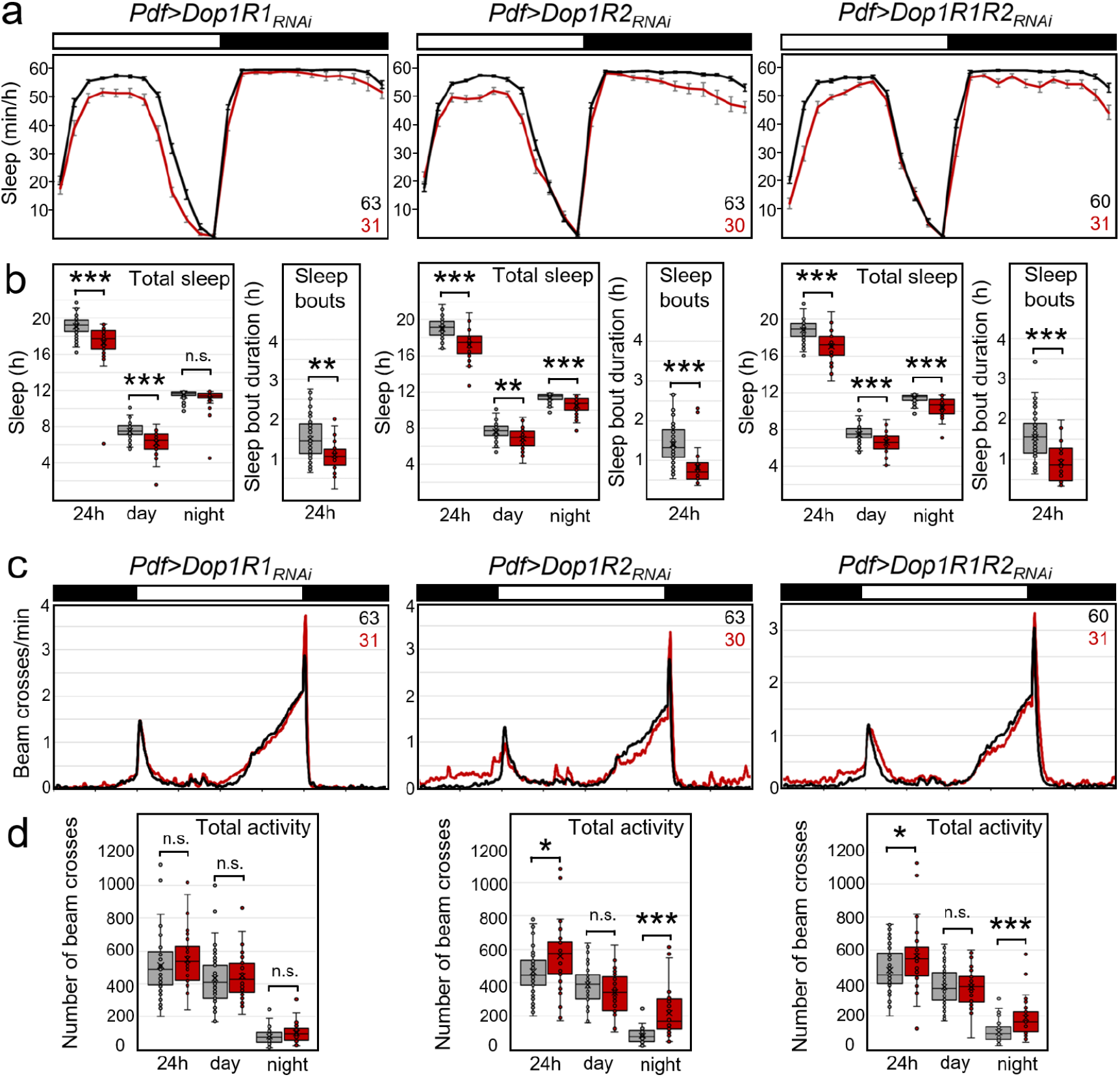
Sleep and activity in PDF-neuron specific dopamine-receptor knockdown flies. Dop1R1, Dop1R2 or both were knocked down using *Pdf*-*Gal4*. (**a**) Average daily sleep profiles of experimental flies (red, *Pdf>Dop1R_RNAi_*) and respective *Gal4* and *UAS* controls (pooled in black; both controls showed significantly more sleep than the flies with dopamine receptor knockdown; therefore they were pooled). (**b**) Box Plots of sleep parameters as shown in Fig. 6. Flies showed significantly less total sleep and shorter sleep bouts, when either Dop1R1 or Dop1R2 or both were knocked down in the PDF-neurons. Knockdown of Dop1R1 decreased daytime sleep, whereas knockdown of Dop1R2 and simultaneous knockdown of both receptors decreased day- and night-time sleep. (**c**) Average activity profiles of the same flies that are depicted in **a**. The flies with down-regulated Dop1R2 receptor were more active than the controls. (**d**) Box Plots of total activity during the entire 24 h period, during the day and the night. Significant differences are indicated by asterisks (* p<0.05; ** p<0.01; *** p<0.001). The numbers of tested flies are indicated in (**a**) and (**c**).

To assess possible developmental effects of Dop1R1 or Dop2R1 knockdown on the PDF neurons, we repeated our LN_v_ knockdown experiments using GeneSwitch (GS) (Depetris-Chauvin et al., 2011). Feeding flies the progesterone derivative RU (dissolved in ethanol) only during adulthood restricted the expression of RNAi constructs to the adult stage. We used two types of controls. (1) *Pdf-GS>uas-Dop1Rx* fed with ethanol alone served as controls for *Pdf-GS>uas-Dop1Rx* flies fed with RU (Fig. 9). (2) *Pdf-GS* and *uas-Dop1Rx* flies, in which the dopamine receptors were not down-regulated and which were fed either with ethanol alone or with RU, served as controls for the effect of RU (Fig. 8). In the latter, we did not find any systematic difference in activity and sleep between the RU and ethanol-fed flies (Fig. 8). Only in *Pdf-GS* controls did we find that nocturnal activity was significantly decreased during the last few hours of the night after feeding RU. In the experimental animals (with dopamine knockdown), the differences between controls and permanent Dop1R2-knockdown during the day disappeared when this receptor knocked-down conditionally, suggesting that these were caused by developmental effects. Nevertheless, the significant reduction in daytime sleep after Dop1R1 knockdown and the reduction of night sleep after Dop1R2 knockdown persisted (Fig. 9a, b). Furthermore, the conditional down-regulation of dopamine receptors increased activity during the day and the night (Fig. 9c, d). Since the effects of conditional dopamine receptor down-regulation were in the same direction as the constitutive receptor down-regulation and in the opposite direction of RU feeding (Fig. 8) in *Pdf-GS* controls, we conclude that these are specific and indeed caused by down-regulation of the dopamine receptors in the PDF neurons.

**Figure 8.**
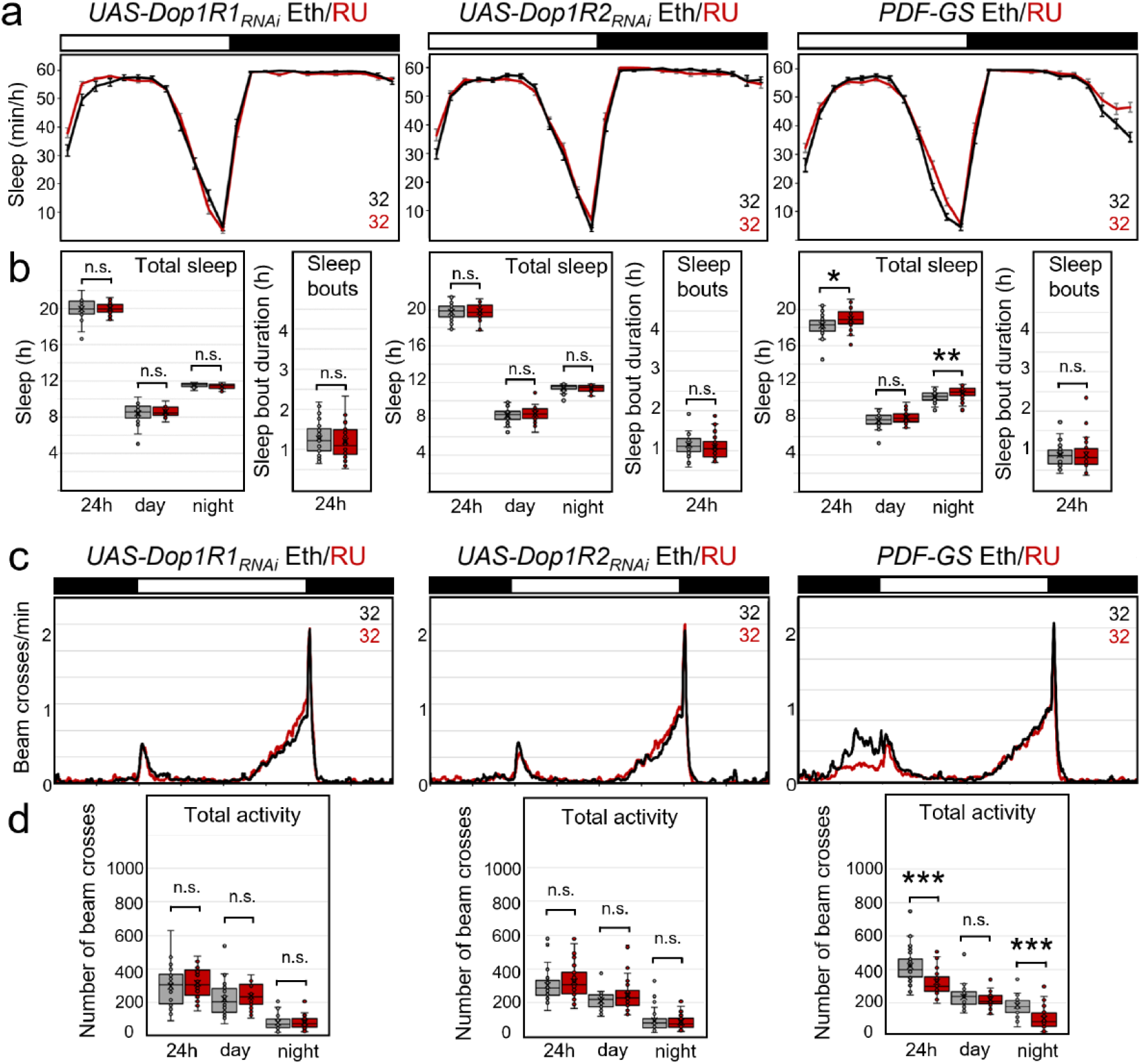
Sleep and activity in control flies fed with RU dissolved in ethanol or only with ethanol. (**a**) Average daily sleep profiles of flies fed with RU in ethanol (red) and flies fed only with ethanol (black). (**b**) Box Plots of sleep parameters. (**c**) Average activity profiles of the same flies that are depicted in **a**. (**d**) Box Plots of total activity during the entire 24 h period, during the day and the night. Significant differences are indicated by asterisks (* p<0.05; ** p<0.01; *** p<0.001). Feeding of RU affected sleep and activity marginally. Only *Pdf-Gal4* flies fed with RU slept significantly more and were less active in the night than flies fed only with ethanol. The numbers of tested flies are indicated in (**a**) and (**c**).

**Figure 9.**
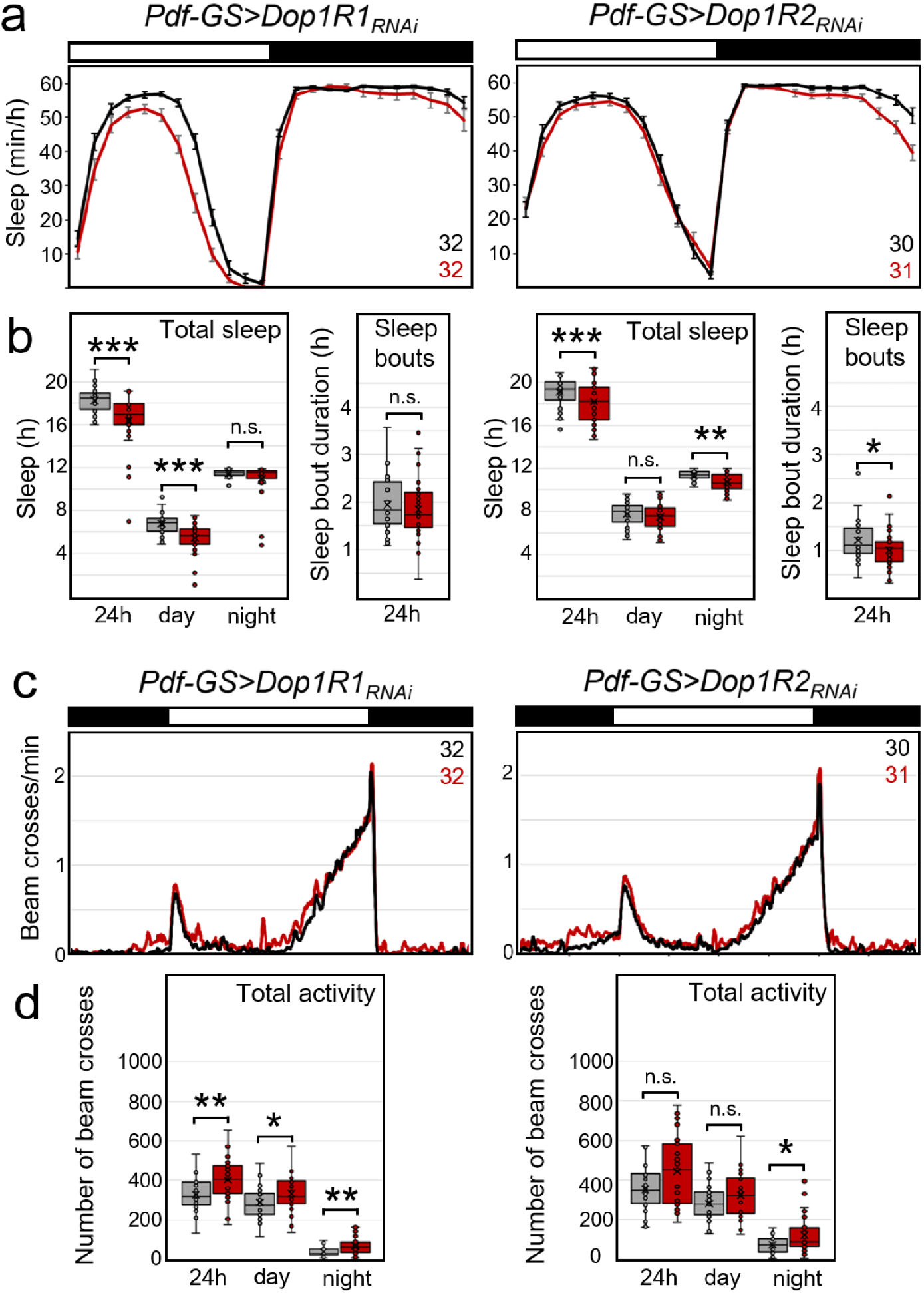
Sleep and activity in flies with conditional dopamine-receptor knockdown in the PDF-neurons (with *Pdf-GS*). (**a**) Average daily sleep profiles of experimental flies (red, *Pdf-GS>Dop1R_RNAi_* fed with RU in ethanol) and control flies (black, *Pdf-GS>Dop1R_RNAi_* fed with ethanol). (**b**) Box Plots of sleep parameters. Flies showed significantly less total sleep when either Dop1R1 or Dop1R2 or both were knocked down in the PDF-neurons. Knockdown of Dop1R1 decreased daytime sleep, whereas knockdown of Dop1R2 decreased nighttime sleep. Sleep bouts were only significantly affected after knockdown of the Dop1R2 receptor. (**c**) Average activity profiles of the same flies that are depicted in **a**. The flies with down-regulated dopamine receptors were generally more active than the controls. (**d**) Box Plots of total activity during the entire 24 h period, during the day and the night. Significant differences are indicated by asterisks (* p<0.05; ** p<0.01; *** p<0.001). The numbers of tested flies are indicated in (**a**) and (**c**).

We observed a highly significant period-lengthening effect of RU application in *Pdf-GS* controls and all the crosses with the *Pdf-GS* strain (Table 1), which has been reported in the past (Depetris-Chauvin et al., 2011; Frenkel et al., 2017). Therefore, we conclude that conditional dopamine receptor down-regulation itself does not affect the free-running period, which is in line with the results obtained via permanent dopamine receptor knockdown.

### Dopamine depolarizes the l-LN_v_s via Dop1R1, but does not increase their firing rate

When observed electrophysiologically using whole-cell patch clamp, the l-LN_v_s fire spontaneous action potentials in bursting or tonic modes (e.g. Cao and Nitabach, 2008; Sheeba et al., 2008b; Depetris-Chauvin et al., 2011; Fogle et al., 2011; Muraro and Ceriani, 2015). As reported previously, when whole-cell patch clamp recordings are performed in the morning and established rapidly after brain dissection (Muraro and Ceriani, 2015), all l-LN_v_s fire action potentials in the bursting mode (Fig. 10). To further explore the role of dopamine on the physiology of l-LN_v_s, we bath-applied dopamine across control l-LN_v_s (Fig. 10a), and in l-LN_v_s in which Dop1R1 (Fig. 10b) or Dop1R2 (Fig. 10c) had been down-regulated using RNAi constructs driven by the *Pdf*-*Gal4*. Control and Dop1R2_RNAi_ l-LN_v_s displayed robust depolarizations upon 1 mM dopamine application (Fig. 10a, c, and d). In contrast, we observed significantly reduced dopamine induced depolarization when Dop1R1 expression was down-regulated (Fig. 10b and d). This result is consistent with cAMP imaging experiments (Fig. 4) and supports the hypothesis that dopamine responses in l-LN_v_s are mainly mediated by the Dop1R1 receptor. Although we observed a small trend toward a decrease in firing rate upon dopamine application, this was not statistically significant (Fig. 11). These results suggest that, in l-LN_v_s, dopamine plays a modulatory role as it depolarizes the membrane without significantly changing the firing rate. Thus, dopamine might make the l-LN_v_s more sensitive to excitatory inputs.

**Figure 10.**
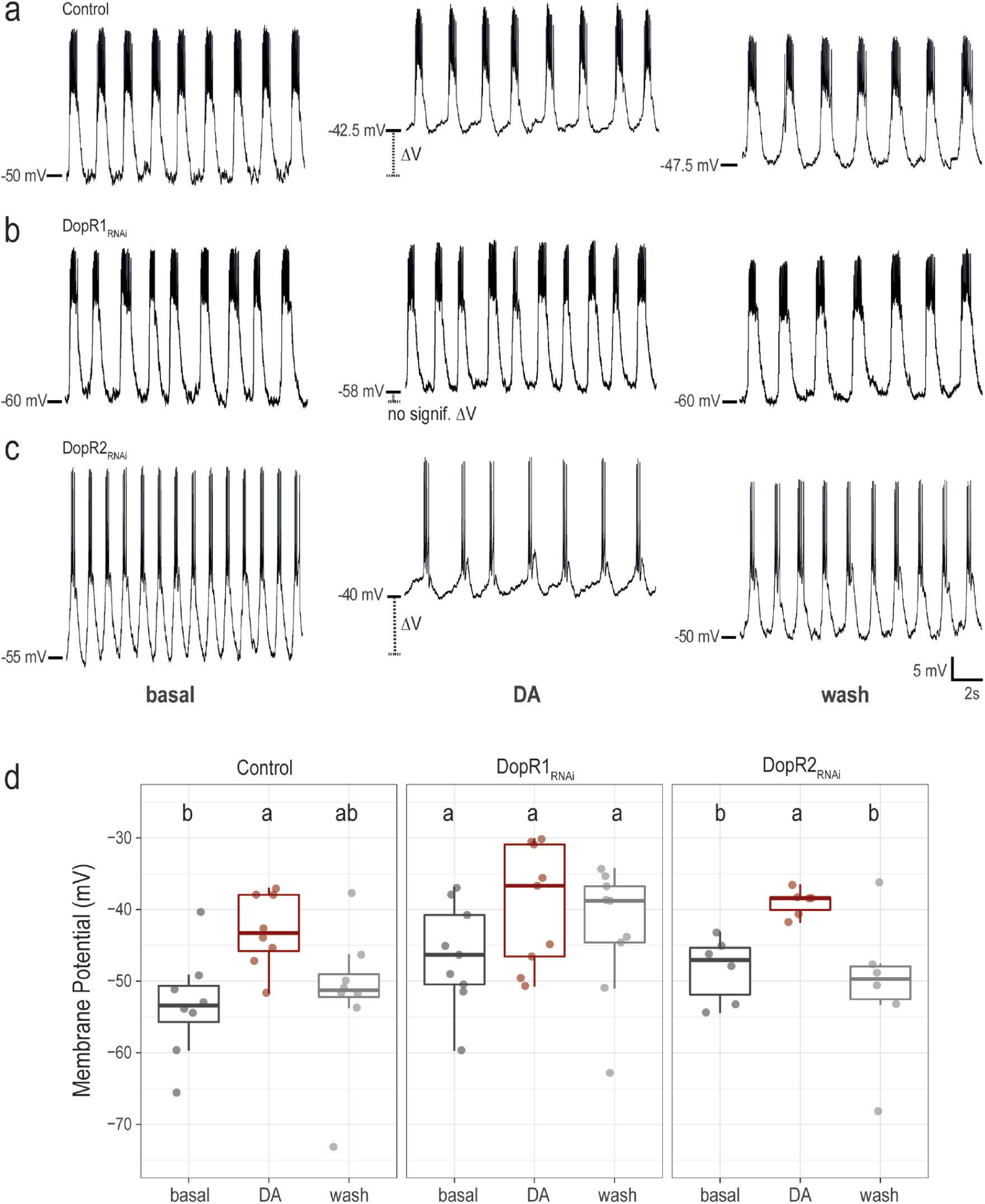
Dop1R1 receptor mediates l-LN_v_ responses to dopamine. (**a**), (**b**), (**c**). Representative traces of whole-cell patch clamp recordings during basal conditions (perfusion of external saline, left panels), DA (perfusion of 1 mM dopamine, middle panels) and wash out (perfusion of external saline, right panels). (**a**) Control group, *Pdf-Gal4,UAS-dicer*;*pdf* Red>+. (**b**), *Dop1R1_RNAi_* group, *Pdf-Gal4,UAS-dicer2;pdf* Red>*UAS-Dop1R1_RNAi_*. (**c**), *Pdf-Gal4,UAS-dicer2;pdf* Red>*UAS-Dop1R2_RNAi_*. (**d**). Boxplots showing the value of membrane potential in mV for the same genotypes in each condition (basal, DA, wash). Kruskal-Wallis non-parametric test with Bonferroni correction was applied for statistical analysis. The alpha parameter was 0.05. Different letters indicate significant differences. Control, n=8. DopR1_RNAi_, n=9. Dop1R2_RNAi_, n=6.

**Figure 11.**
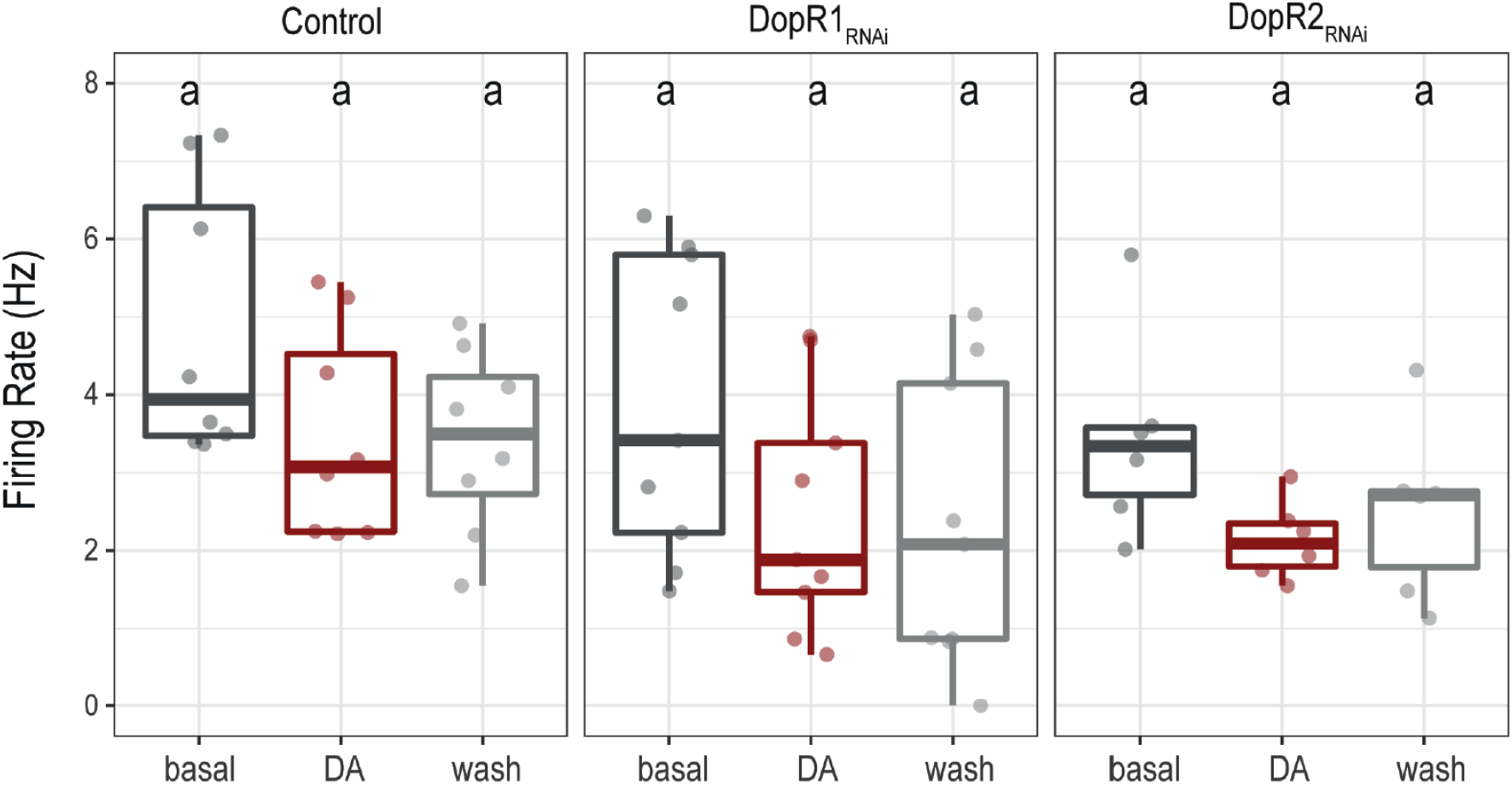
Boxplots showing the value of firing rate (number of action potentials per second) obtained in whole-cell patch clamp configuration under three different conditions (basal, Dopamine, wash) for Pdf-Gal4,UAS-dicer;pdf Red>+ (Control, left panel), DopR1RNAi group, Pdf-Gal4,UAS-dicer;pdf Red>UAS-Dop1R1RNAi (DopR1RNAi, middle panel), Pdf-Gal4,UAS-dicer;pdf Red>UAS-Dop1R2RNAi (Dop1R2RNAi, right panel). Kruskal-Wallis non-parametric test with Bonferroni correction was applied for statistical analysis. The alpha parameter was 0.05. No statistically significant differences were found (same letter indicate no significant differences). Control, n=8. Dop1R1_RNAi_, n=9. Dop1R2_RNAi_, n=6.

## Discussion

### All tested clock neurons respond to dopamine

Here we show that dopamine acts broadly on the neurons of the *Drosophila* clock network that have neurites in the AME, a neuropil that is invaded by presynaptic terminals of dopaminergic neurons. All of these clock neurons responded to dopamine with increases in cAMP. The responses of the l-LN_v_s and DN_1_s were almost completely blocked by downregulation of Dop1R1 receptors but not significantly by down-regulation of Dop1R2 receptors, whereas the responses of some LN_d_s were blocked by down-regulation of Dop1R1 and others by down-regulation of Dop1R2 receptors. Dopamine responses of all LN_d_ cells were eliminated by simultaneous down-regulation of both receptors. This indicates that the LN_d_s employ different activating dopamine receptors.

Since the electrophysiological and cAMP responses of the l-LN_v_s were not blocked by down-regulating Dop1R2 receptors we conclude that these neurons employ only Dop1R1 receptors. Unfortunately, we could not assess the nature of the Dop1R receptors in the s-LN_v_s, but we hypothesize that these employ Dop1R2 receptors for the following reason: the downregulation of Dop1R2 receptors in the s-LN_v_s and l-LN_v_s significantly reduces the flies’ nighttime sleep. Since the l-LN_v_s appear not to utilize Dop1R2 receptors this effect is most likely mediated by the s-LN_v_s.

### Dopamine signaling on the s-LN_v_s appears to promote sleep

Multiple lines of evidence are consistent with a wake promoting role for the s-LN_v_s (e.g. Liang et al., 2019). We were therefore surprised to find that the knockdown of the excitatory dopamine receptor Dop1R2 produce decreases in nighttime sleep. We note here that the s-LN_v_s have been shown to promote sleep during the entire day via PDF-signaling to the AllatostatinA (AstA) positive ‘PLP’ neurons (Chen et al., 2016), which were recently shown to be identical with the Lateral Posterior clock neurons (LPNs) (Ni et al., 2019). Optogenetic excitation of the LPNs promotes sleep (Guo et al. 2018) and glutamatergic and AstA neurites provide excitatory inputs on to the sleep promoting dorsal fan-shaped body (Donlea et al., 2011; Liu et al., 2012; 2016; Ueno et al., 2012; Pimentel et al., 2016; Ni et al., 2019). Thus, our results, along with previous work, suggest that: 1) the role of the s-LN_v_s in the control of sleep is more complex than previously acknowledged, 2) dopamine likely increases cAMP levels in the s-LN_v_s via Dop1R2, 3) the s-LN_v_s excite the sleep promoting LPNs, which subsequently activate the dorsal fan-shaped body neurons leading to sleep. Thus, down-regulation of Dop1R2 receptors in the s-LN_v_s would therefore be predicted to reduce sleep, which fits to our observations and is consistent with the literature. However, we also must acknowledge the possibility that the s-LN_v_s might promote both sleep and wakefulness at different times. Recent work on the DN_1p_ class of clock neurons showed that the temporal codes of firing in these cells shape sleep (Tabuchi et al. 2018), suggesting that some clock neurons can switch between sleep and wake promoting modes through changes in their patterns of firing. The same may prove true of the s-LN_v_s.

### Dopamine signaling on the l-LN_v_s is not wake-promoting

The l-LN_v_s were reported to be strongly wake-promoting (Sheeba et al., 2008a; Chung et al., 2009; Shang et al., 2011), but it was not clear if dopamine-signaling was responsible this effect. Here, we could not detect wake-promoting effects of dopamine signaling on the PDF neurons. In contrast, down-regulation of the excitatory Dop1R1 and Dop1R2 receptors in these neurons (along with the s-LN_v_s) slightly increased wakefulness. Night-sleep decreased after knockdown of Dop1R2 receptors, while day-sleep decreased after knockdown of Dop1R1 receptors. Our physiological observations make it clear that and Dop1R1 receptors are expressed by the l-LN_v_s. This evidently speaks against a wake-promoting role of dopamine signaling to l-LN_v_s.

The present study supports the findings of Ueno et al. (2012) who found that the ablation of the l-LN_v_s did not eliminate the strong arousal effects of dopamine, thereby suggesting that dopamine does not drive the wake-promoting role of the l-LN_v_s. In fact, our results suggest a moderate sleep-promoting effect of dopamine signaling on the l-LN_v_s, despite of the fact that dopamine depolarizes the l-LN_v_s, potentially making them more excitable. Glutamate, GABA, and histamine inhibit the l-LN_v_s (Cao and Nitabach, 2008; Schlichting et al., 2016). While GABAergic inputs to l-LN_v_s have a clear role in the promotion of sleep (Agosto et al., 2008; Parisky et al., 2008; Chung et al., 2009; Gmeiner et al., 2013), such a role has not yet been demonstrated for histamine or glutamate. Other putative silencing neuromodulators of the l-LN_v_s are glycine (Frenkel et al., 2017) and serotonin (Yuan et al., 2005, 2006), but how these different signals interact to regulate the l-LN_v_s’ command over wakefulness is still an open question.

Our study does not call into question the wake-promoting role of the l-LN_v_s. The ablation of the l-LN_v_s increases sleep, which demonstrates that their wake-promoting influence exceeds their sleep-promoting one (Chung et al., 2009). Furthermore, the l-LN_v_s are electrically the most active during the day when the flies are awake (Sheeba et al., 2008b; Shang et al., 2011) and the electrical hyperexcitation of the l-LN_v_s increases activity at night and disrupts nocturnal sleep (Sheeba et al., 2008a). Thus, the l-LN_v_s are firing during the day, thereby promoting daytime wakefulness, and their firing is decreased at night when flies maintain their deepest sleep. The wake promoting neuromodulators octopamine and acetylcholine act on l-LN_v_s (Kula-Eversole et al., 2010; Muraro and Ceriani, 2015). But the result described above, lead to the surprising conclusion that dopamine does not act wake-promoting neuromodulator of the l-LN_v_s.

In any case, the sleep-promoting role of dopamine via the l-LN_v_s is moderate when compared to the sleep-promoting effects of the fan-shaped body neurons that lack dopaminergic input (Liu et al., 2012; Ueno et al., 2012). Thus, dopamine signaling via the fan-shaped body has a stronger impact on sleep than dopamine signaling via the l-LN_v_s or the s-LN_v_s. The precise role played by dopaminergic inputs to l-LN_v_s and their modulatory effect on the integration of the multiple excitatory and inhibitory afferences received by these important arousal neurons awaits further research.

### Dopamine has different effects on the fan-shaped body neurons and the PDF neurons

Dop1R1 and Dop1R2 receptors have already been implicated in the control of sleep in previous studies. Lebestky et al. (2009) showed that the rescue of Dop1R1 receptors in the l-LN_v_s of Dop1R1 mutants can partially rescue the flies’ normal sleep pattern, which fits our observation that the l-LN_v_s utilize Dop1R1 receptors. Liu et al. (2012) and Ueno et al. (2012) showed that dopaminergic neurons signal via Dop1R1 receptors on neurons in the fan-shaped body whereas Pimentel et al. (2016) demonstrated a role of Dop1R2 receptors in the fan-shaped body. Here we suggest that dopamine signals via Dop1R2 receptors on the s-LN_v_s. Although the PDF neurons and the fan-shaped body neurons respond to dopamine via the same activating receptors and in both cases via an increase in cAMP levels, the electrical responses of the neurons to dopamine appear to be different.

In the fan-shaped body neurons, the increase of cAMP leads to an upregulation of the voltage-independent leak current K^+^ channel “Sandman” and its translocation to the plasma membrane (Pimentel et al., 2016). Consequently, the fan-shaped body neurons switch to long-lasting hyperpolarization (OFF state), which keeps the fruit flies awake. The Rho-GTPase-activating protein Crossveinless-c locks the fan-shaped body neurons in the OFF state (Donlea et al., 2014) until unknown mechanisms flip the neurons back to the ON state. Thus, Dop1R1/2 receptors silence neurons in the fan-shaped body via the increase of cAMP levels (Liu et al., 2012; Ueno et al., 2012; Pimentel et al., 2016).

Our results indicate a very different effect of Dop1R1 receptor signaling in the l-LN_v_s. The neurons depolarized in response to dopamine and this effect was blocked after knockdown of Dop1R1 receptors. Thus, dopamine excites the l-LN_v_s as predicted, but does not increase their firing rate. The main effect of dopamine perfusion in our *ex-vivo* preparation was a robust and reversible depolarization of the membrane, which should make l-LN_v_s more sensitive to excitatory inputs. Thus, the effect of dopamine on the l-LN_v_s may be contextdependent. Lebestky et al. (2009) aroused the flies by repetitive air puffs and found that dopamine reduced the flies’ hyperactivity in response to this excitation, while it increased spontaneous nocturnal activity. Both effects were mediated via Dop1R1 receptors. Although Lebestky et al. (2009) traced the dopamine effects on startle-induced hyperactivity to the central complex, we cannot exclude that similar mechanisms work in the l-LN_v_s. Therefore, it will be most interesting to study the effects of Dop1R1 receptor knock-down in the l-LN_v_s on sleep and activity of flies in the context of stimulus-induces arousal, to test not only the role of dopaminergic inputs to l-LN_v_s in the context of basal sleep-wake activity, but also in the context of environmentally stimulated arousal or in the presence of challenges to the sleep homeostat, such as in the generation of a sleep rebound phenomenon after a night of sleep deprivation.

In summary, dopamine appears to have different modulatory effects on the fan-shaped body neurons and the PDF neurons – inhibiting the former and exciting the latter. In both cases, dopamine signaling increases sleep, though in different ways and to different degrees. Dopamine signaling to the fan-shaped body is strongly sleep promoting, while dopamine signaling to the PDF neurons is weakly sleep promoting and, in case of the l-LN_v_s, perhaps dependent on the arousal state of the flies.

## Declarations

### Author contributions

FF-C performed and analyzed the patch-clamp recordings, CH-L performed and analyzed the cAMP imaging and the behavioral experiments with permanent dopamine receptor knockdown, AP performed the behavioral experiments with conditional dopamine receptor knockdown, NL did the statistical analysis, MH and MS performed the histology, OTS, NIM and CH-F designed the study and supervised the experiments, CH-F analyzed the behavioral experiments with conditional dopamine receptor knockdown and wrote the paper with contributions from CH-L, OTS, NIM and MS.

## Acknowledgements and Funding

This study was supported by the German Research Foundation (DFG; grant Fo207/14-1 and PA3241/2-1) to CHF and MS respectively, the Agencia Nacional de Promoción Científica y Tecnológica of Argentina (grant PICT-2015-2557) to NIM, and FOCEM-Mercosur (COF 03/11) to IBioBA, and by a National Institutes of Health NINDS grant (R01NS077933) and an NSF IOS grant (1354046) to OTS. We thank Serge Birman for providing the TH-Gal4 line and for profound discussions on dopamine effects, Jan Marek Ache for valuable discussion and editing of the manuscript, Indra Hering for help with the sleep experiments and Barbara Mühlbauer for general excellent assistance.

## Ethics approval

Not applicable

## References

Agosto J, Choi JC, Parisky KM, Stilwell G, Rosbash M, Griffith LC (2008) Modulation of GABAA receptor desensitization uncouples sleep onset and maintenance in *Drosophila*. Nat Neurosci 11:354–359. https://doi.org/10.1038/nn2046

Andretic R, van Swinderen B, Greenspan RJ (2005) Dopaminergic modulation of arousal in *Drosophila*. Curr Biol 15:1165–1175. https://doi.org/10.1016/j.cub.2005.05.025

Birman S (2005) Arousal mechanisms: speedy flies don’t sleep at night. Curr Biol 15(16):R511–513. https://doi.org/10.1016/j.cub.2005.06.032

Cao G, Nitabach, MN (2008) Circadian control of membrane excitability in *Drosophila melanogaster* lateral ventral clock neurons. J Neurosci 28:6493–6501. https://doi.org/10.1523/JNEUROSCI.1503-08.2008

Cavey M, Collins B, Bertet C, Blau J (2016) Circadian rhythms in neuronal activity propagate through output circuits. Nat Neurosci 19:587–595. https://doi.org/10.1038/nn.4263

Chen J, Reiher W, Hermann-Luibl C, Sellami A, Cognigni P, Kondo S, Helfrich-Förster C, Veenstra JA, Wegener C (2016) Allatostatin A signalling in *Drosophila* regulates feeding and sleep and is modulated by PDF. PLoS Genetics 12(9):e1006346. https://doi.org/10.1371/journal.pgen.1006492

Chung BY, Kilman VL, Keath JR, Pitman JL, Allada R (2009) The GABAA receptor RDL acts in peptidergic PDF neurons to promote sleep in *Drosophila*. Curr Biol 19(5):386–390. https://doi.org/10.1016/j.cub.2009.01.040

Cirelli C (2009) The genetic and molecular regulation of sleep: from fruit flies to humans. Nat Rev 10:549–560. https://doi.org/10.1038/nrn2683

Crocker A, Shahidullah M, Levitan IB, Sehgal A (2010) Identification of a neural circuit that underlies the effects of octopamine on sleep: wake behavior. Neuron 65:670–681.. https://doi.org/10.1016/j.neuron.2010.01.032

Depetris-Chauvin A, Berni J, Aranovich EJ, Muraro NI, Beckwith EJ, Ceriani MF (2011) Adultspecific electrical silencing of pacemaker neurons uncouples molecular clock from circadian outputs. Curr Biol 21, 1783–1793. https://doi.org/10.1016/j.cub.2011.09.027

Donlea JM, Ramanan N, Shaw PJ (2009) Use-dependent plasticity in clock neurons regulates sleep need in *Drosophila*. Science 324:105–108. https://doi.org/10.1126/science.1166657

Donlea JM, Thimgan MS, Suzuki L, Gottschalk L, Shaw PJ (2011) Inducing sleep by remote control facilitates memory consolidation in *Drosophila*. Science 332:1571–1576. https://doi.org/10.1126/science.1202249

Donlea JM, Pimentel D, Miesenböck G (2014) Neuronal machinery of sleep homeostasis in *Drosophila*. Neuron 81:860–872. https://doi.org/10.1016/j.neuron.2013.12.013

Donlea JM, Pimentel D, Talbot CB, Kempf A, Omoto JJ, Hartenstein V, Miesenböck G (2018) Recurrent Circuitry for Balancing Sleep Need and Sleep. Neuron 97:378–389. https://doi.org/10.1016/j.neuron.2017.12.016

Dubowy C, Sehgal A (2017) Circadian Rhythms and sleep in *Drosophila melanogaster*. Genetics 205(4):1373–1397. https://doi.org/10.1534/genetics.115.185157

Feinberg EH, Vanhoven MK, Bendesky A, Wang G, Fetter RD, Shen K, Bargmann CI (2008) GFP Reconstitution Across Synaptic Partners (GRASP) defines cell contacts and synapses in living nervous systems. Neuron 57:353–363. https://doi.org/10.1016/j.neuron.2007.11.030

Fogle KJ, Parson KG, Dahm NA, Holmes TC (2011) CRYPTOCHROME is a blue-light sensor that regulates neuronal firing rate. Science 331:1409–1413. https://doi.org/10.1126/science.1199702

Foltenyi K, Greenspan RJ, Newport JW (2007) Activation of EGFR and ERK by rhomboid signaling regulates the consolidation and maintenance of sleep in *Drosophila*. Nat Neurosci 10:1160–1167. https://doi.org/10.1038/nn1957

Frenkel L, Muraro NI, Gonzáles ANB, Márcora MS, Bernabó G, Hermann-Luibl C, Romero JI, Helfrich-Förster C, Castano EM, Marino-Busjle C, Calvo DJ, Ceriani MF (2017) Organization of circadian behavior relies on glycinergic transmission. Cell Reports 19:72–85. https://doi.org/10.1016/j.celrep.2017.03.034

Friggi-Grelin F, Coulom H, Meller M, Gomez D, Hirsh J, Birman S (2002) Targeted gene expression in *Drosophila* dopaminergic cells using regulatory sequences from tyrosine hydroxylase. J Neurobiol 54:618–627. https://doi.org/10.1002/neu.10185

Glaser W (1978) Varianzanalyse. Gustav Fischer Verlag. Stuttgart, New York.

Gmeiner F, Kołodziejczyk A, Yoshii T, Rieger D, Nässel DR, Helfrich-Förster C (2013) GABAB receptors play an essential role in maintaining sleep during the second half of the night in *Drosophila melanogaster*. J Exp Biol 216(20):3837–3843. https://doi.org/10.1242/jeb.085563

Grima B, Chélot E, Xia R, Rouyer F (2004) Morning and Evening Peaks of Activity Rely on Different Clock Neurons of the Drosophila Brain. Nature 431(7010):869–73. https://doi.org/10.1038/nature02935

Gummadova JO, Coutts GA, Glossop NR (2009) Analysis of the *Drosophila* Clock promoter reveals heterogeneity in expression between subgroups of central oscillator cells and identifies a novel enhancer region. J Biol Rhythms 24(5):353–367. https://doi.org/10.1177/0748730409343890

Guo F, Yu J, Jung HJ, Abruzzi KC, Luo W, Griffith LC, Rosbash M (2016) Circadian neuron feedback controls the *Drosophila* sleep-activity profile. Nature 536, 292–297. https://doi.org/10.1038/nature19097

Guo F, Holla M, Díaz MM, Rosbash M (2018) A Circadian Output Circuit Controls Sleep-Wake Arousal in Drosophila. Neuron 100, 1–12. https://doi.org/10.1016/j.neuron.2018.09.002

Hamasaka Y, Nässel DR (2006) Mapping of serotonin, dopamine, and histamine in relation to different clock neurons in the brain of *Drosophila*. J Comp Neurol 494:314–330. https://doi.org/10.1002/cne.20807

Helfrich-Förster C (1995) The *period* clock gene is expressed in central nervous system neurons which also produce a neuropeptide that reveals the projections of circadian pacemaker cells within the brain of *Drosophila melanogaster*. Proc Natl Acad Sci USA 92:612–616. https://doi.org/10.1073/pnas.92.2.612

Helfrich-Förster C (2018) Sleep in insects. Ann Rev Entomol 63:69–86. https://doi.org/10.1146/annurev-ento-020117-043201.

Helfrich-Förster C, Shafer OT, Wülbeck C, Grieshaber E, Rieger D, Taghert P (2007) Development and morphology of the clock-gene-expressing Lateral Neurons of *Drosophila melanogaster*. J Comp Neurol 500:47–70. https://doi.org/10.1002/cne.21146

Hendricks JC, Finn SM, Panckeri KA, Chavkin J, Williams JA, Sehgal A, Pack AI (2000) Rest in *Drosophila* is a sleep-like state. Neuron 25:129–318. https://doi.org/10.1016/S0896-6273(00)80877-6

Hermann-Luibl C, Yoshii T, Senthilan PR, Dircksen H, Helfrich-Förster C (2014) The Ion Transport Peptide is a new functional clock neuropeptide in the fruit fly *Drosophila melanogaster*. J Neurosci 34(29), 9522–9536. https://doi.org/10.1016/j.cois.2014.11.003

Joiner WJ, Crocker A, White BH, Sehgal A (2006) Sleep in *Drosophila* is regulated by adult mushroom bodies. Nature 441:757–760. https://doi.org/10.1038/nature04811

King AN, Sehgal A (2020) Molecular and circuit mechanisms mediating circadian clock output in the *Drosophila* brain. Eur J Neurosci 51(1):268–281. https://doi.org/10.1111/ejn.14092

Kula-Eversole E, Nagoshi E, Shang Y, Rodriguez J, Allada R (2010) Surprising gene expression patterns within and between PDF-containing circadian neurons in *Drosophila*. PNAS 107:13497–13502. https://doi.org/10.1073/pnas.1002081107

Kume K, Kume S, Park SK, Hirsh J, Jackson FR (2005) Dopamine is a regulator of arousal in the fruit fly. J Neurosci 25:7377–7384. https://doi.org/10.1523/JNEUROSCI.2048-05.2005

Lebestky T, Chang JS, Dankert H, Zelnik L, Kim YC, Han KA, Wolf FW, Perona P, Anderson DJ (2009) Two different forms of arousal in *Drosophila* are oppositely regulated by the dopamine D1 receptor ortholog DopR via distinct neural circuits. Neuron 64:522–536. https://doi.org/10.1016/j.neuron.2009.09.031

Liang X, Ho MCW, Zhang Y, Li Y, Wu MN, Holy TE, Taghert PH (2019). Morning and Evening Circadian Pacemakers Independently Drive Premotor Centers via a Specific Dopamine Relay. Neuron 102(4):843–857. https://doi.org/10.1016/j.neuron.2019.03.028

Lima SQ, Miesenböck G (2005) Remote control of behavior through genetically targeted photostimulation of neurons. Cell 121(1):141–152. https://doi.org/10.1016/j.cell.2005.02.004

Liu S, Lamaze A, Liu Q, Tabuchi M, Yang Y, Fowler M, Bharadwaj R, Zhang J, Bedont J, Blackshaw S, Lloyd TE, Montell C, Sehgal A, Koh K, Wu MN (2014) WIDE AWAKE mediates the circadian timing of sleep onset. Neuron 82:151–166. https://doi.org/10.1016/j.neuron.2014.01.040

Liu S, Liu Q, Tabuchi M, Wu MN (2016). Sleep drive is encoded by neural plastic changes in a dedicated circuit. Cell 165:1347–1360. https://doi.org/10.1016/j.cell.2016.04.013

Liu Q, Liu S, Kodama MR, Driscoll MR, Wu MN (2012) Two dopaminergic neurons signal to the dorsal fan-shaped body to promote wakefulness in *Drosophila*. Curr Biol 22:2114–2123. https://doi.org/10.1016/j.cub.2012.09.008

Muraro N, Ceriani MF (2015) Acetylcholine from visual circuits modulates the activity of arousal neurons in *Drosophila*. J Neurosci 35(50):16315–16327. https://doi.org/10.1523/JNEUROSCI.1571-15.2015

Ni JD, Gurav AS, Liu W, Ogunmowo TH, Hackbart H, Elsheikh A, Verdegaal AA, Montell C (2019) Differential regulation of the *Drosophila* sleep homeostat by circadian and arousal inputs. Elife 8:e40487. https://doi.org/10.7554/eLife.40487

Nikolaev VO, Bünemann M, Hein L, Hannawacker A, Lohse MJ (2004) Novel single chain cAMP sensors for receptor-induced signal propagation. J Biol Chem 279(36):37215–37218. https://doi.org/10.1074/jbc.C400302200

Nitz DA, van Swinderen B, Tononi G, Greenspan RJ (2002) Electrophysiological correlates of rest and activity in *Drosophila melanogaster*. Curr Biol 12:1934–1940. https://doi.org/10.1016/s0960-9822(02)01300-3

Osterwalder T, Yoon KS, White BH, Keshishian H (2001) A conditional tissue-specific transgene expression system using inducible GAL4. Proc Natl Acad Sci USA 98: 12596–12601. https://doi.org/10.1073/pnas.221303298

Parisky KM, Agosto J, Pulver SR, Shang Y, Kuklin E, Hodge JJ, Kang K, Liu X, Garrity PA, Rosbash M, Griffith LC (2008) PDF cells are a GABAresponsive wake-promoting component of the *Drosophila* sleep circuit. Neuron 60:672–682. https://doi.org/10.1016/j.neuron.2008.10.042

Park S, Sonn JY, Oh Y, Lim C, Choe J (2014). SIFamide and SIFamide receptor define a novel neuropeptide signaling to promote sleep in *Drosophila*. Mol. Cells 37:295–301. https://doi.org/10.14348/molcells.2014.2371

Pimentel D, Donlea JM, Talbot CB, Song SM, Thurston AJ, Miesenböck G (2016) Operation of a homeostatic sleep switch. Nature 536:333–337. https://doi.org/10.1038/nature19055

Pitman JL, McGill JJ, Keegan KP, Allada R (2006) A dynamic role for the mushroom bodies in promoting sleep in *Drosophila*. Nature 441:753–756. https://doi.org/10.1038/nature04739

Potdar S, Sheeba V (2018) Wakefulness is promoted during day time by PDFR signalling to dopaminergic neurons in *Drosophila melanogaster*. eNeuro 5(4): 0129–18. https://doi.org/10.1523/ENEURO.0129-18

Renn SC, Park JH, Rosbash M, Hall JC, Taghert PH (1999) A Pdf Neuropeptide Gene Mutation and Ablation of PDF Neurons Each Cause Severe Abnormalities of Behavioral Circadian Rhythms in Drosophila. Cell 99(7):791–802. https://doi.org/10.1016/S0092-8674(00)81676-1 https://doi.org/10.1523/JNEUROSCI.1234-05.2006

Rieger D, Shafer OT, Tomioka K and Helfrich-Förster Ch (2006) Functional Analysis of Circadian Pacemaker Neurons in Drosophila melanogaster. Journal of Neuroscience 26 (9) 2531–2543. https://doi.org/10.1523/JNEUROSCI.1234-05.2006

Riemensperger T, Isabel G, Coulom H, Neuser K, Seugnet L, Kume K, Iché-Torres M, Cassar M, Strauss R, Preat T, Hirsh J, Birman S (2011) Behavioral consequences of dopamine deficiency in the *Drosophila* central nervous system. PNAS 108:834–839. https://doi.org/10.1073/pnas.1010930108

Ruben M, Drapeau MD, Mizrak D, Blau J (2012) A mechanism for circadian control of pacemaker neuron excitability. J Biol Rhythms 27:353–364. https://doi.org/10.1177/0748730412455918

Schlichting, Helfrich-Förster C (2015) Photic entrainment in Drosophila assessed by locomotor activity recordings. Methods Enzymol. 552:105–23. https://doi.org/10.1016/bs.mie.2014.10.017

Schlichting M, Menegazzi P, Lelito KR, Yao Z, Buhl E, Dalla Benetta E, Bahle A, Denike J, Hodge JJL, Helfrich-Förster C, Shafer OT (2016) A neural network underlying circadian entrainment and photoperiodic adjustment of sleep and activity in *Drosophila*. J Neurosci 36:9084–9096. https://doi.org/10.1523/JNEUROSCI.0992-16.2016

Schubert FK, Hagedorn N, Yoshii T, Helfrich-Förster C, Rieger D (2018) Neuroanatomical details of *Drosophila* lateral neurons support their functional role in the circadian system. J Comp Neurol 526(7):1209–1231. https://doi.org/10.1002/cne.24406

Shafer OT, Kim DJ, Dunbar-Yaffe R, Nikolaev VO, Lohse MJ, Taghert PH (2008) Shafer OT, Kim DJ, Dunbar-Yaffe R, Nikolaev VO, Lohse MJ, Taghert PH. Neuron 58(2):223–37. https://doi.org/10.1016/j.neuron.2008.02.018

Shang Y, Griffith LC, Rosbash M (2008) Light-arousal and circadian photoreception circuits intersect at the large PDF cells of the *Drosophila* brain. PNAS 105:19587–1994. https://doi.org/10.1073/pnas.0809577105

Shang Y, Haynes P, Pírez N, Harrington KI, Guo F, Pollack J, Hong P, Griffith LC, Rosbash M (2011) Imaging analysis of clock neurons reveals light buffers the wake-promoting effect of dopamine. Nat Neurosci 14:889–895. https://doi.org/10.1038/nn.2860

Shaw PJ, Cirelli C, Greenspan RJ, Tononi G (2000) Correlates of sleep and waking in *Drosophila melanogaster*. Science 287:1834–1837. https://doi.org/10.10.1126/science.287.5459.1834

Sheeba V, Fogle J, Kaneko M, Rashid S, Chou Y-T (2008a) Large ventral lateral neurons modulate arousal and sleep in *Drosophila*. Curr Biol 18:1537–1545. https://doi.org/10.1016/j.cub.2008.08.033

Sheeba V, Gu H, Sharma VK, O’Dowd DK, Holmes TC (2008b) Circadian-and light-dependent regulation of resting membrane potential and spontaneous action potential firing of Drosophila circadian pacemaker neurons. J Neurophysiol 99:976–988. https://doi.org/10.1152/jn.00930.2007

Stewart, BA, Atwood HL, Renger JJ, Wang J, Wu CF (1994) Improved stability of Drosophila larval neuromuscular preparations in haemolymph-like physiological solutions. Journal of Comparative Physiology 175, 179–191. https://doi.org/10.1007/BF00215114

Stoleru D, Peng Y, Agosto J, Rosbash M (2004) Coupled Oscillators Control Morning and Evening Locomotor Behaviour of Drosophila. Nature 431(7010):862–8. https://doi.org/10.1038/nature02926

Tabuchi M, Monaco JD, Duan G, Bell B, Liu S, Liu Q, Zhang K, Wu MN (2018). Clock-Generated Temporal Codes Determine Synaptic Plasticity to Control Sleep. Cell 175(5):1213–1227. https://doi.org/10.1016/j.cell.2018.09.016

Ueno T, Tomita J, Tanimoto H, Endo K, Ito K, Kume S, Kume K (2012). Identification of a dopamine pathway that regulates sleep and arousal in *Drosophila*. Nat Neurosci 15:1516–1523. https://doi.org/10.1038/nn.3238

Van Swinderen B, Nitz DA, Greenspan RJ (2004) Uncoupling of brain activity from movement defines arousal States in *Drosophila*. Curr Biol 14(2):81–87. https://doi.org/10.1016/j.cub.2003.12.057

Wang J, Ma X, Yang JS, Zheng X, Zugates CT, Lee CHJ, Lee T (2004) Transmembrane/juxtamembrane domain-dependent Dscam distribution and function during mushroom body neuronal morphogenesis. Neuron 43:663–672. https://doi.org/10.1016/j.neuron.2004.06.033

Wu MN, Koh K, Yue Z, Joiner WJ, Sehgal A (2008) A genetic screen for sleep and circadian mutants reveals mechanisms underlying regulation of sleep in *Drosophila*. Sleep 31:465–472.

Yuan Q, Lin F, Zeng X, Sehgal A (2005) Serotonin modulates circadian entrainment in *Drosophila*. Neuron 47(1):115–127. https://doi.org/10.1093/sleep/31.4.465

Yuan Q, Joiner WJ, Sehgal A (2006) A sleep-promoting role for the *Drosophila* serotonin receptor 1A. Curr Biol 16:1051–1062. https://doi.org/10.1016/j.cub.2006.04.032

